# Octopaminergic neurons have multiple targets in *Drosophila* larval mushroom body calyx and regulate behavioral odor discrimination

**DOI:** 10.1101/295659

**Authors:** J Y Hilary Wong, Bo Angela Wan, Tom Bland, Marcella Montagnese, Alex McLachlan, Cahir J O’Kane, Shuo Wei Zhang, Liria M Masuda-Nakagawa

## Abstract

Discrimination of sensory signals is essential for an organism to form and retrieve memories of relevance in a given behavioural context. Sensory representations are modified dynamically by changes in behavioral state, facilitating context-dependent selection of behavior, through signals carried by noradrenergic input in mammals, or octopamine (OA) in insects. To understand the circuit mechanisms of this signaling, we characterized the function of two OA neurons, sVUM1 neurons, that originate in the subesophageal zone (SEZ) and target the input region of the memory center, the mushroom body (MB) calyx, in larval *Drosophila*. We find that sVUM1 neurons target multiple neurons, including olfactory projection neurons (PNs), the inhibitory neuron APL, and a pair of extrinsic output neurons, but relatively few mushroom body intrinsic neurons, Kenyon cells. PN terminals carried the OA receptor Oamb, a *Drosophila* α1-adrenergic receptor ortholog. Using an odor discrimination learning paradigm, we showed that optogenetic activation of OA neurons compromised discrimination of similar odors but not learning ability. Our results suggest that sVUM1 neurons modify odor representations via multiple extrinsic inputs at the sensory input area to the MB olfactory learning circuit.

## Introduction

Behavioral choices depend on discrimination among “sensory objects”, which are neural representations of multiple coincident sensory inputs, across a range of sensory modalities. For example, “odor objects” (Gottfried 2009; Wilson and Sullivan 2011; Gire et al. 2013) are represented in sparse ensembles of neurons, that are coincidence detectors of multiple parallel inputs from odor quality channels. This principle is used widely in animals, including in mushroom bodies (MBs), the insect center for associative memory (Masuda-Nakagawa et al. 2005; Honegger et al. 2011), and in the piriform cortex (PCx) of mammals (Stettler and Axel 2009; Davison and Ehlers 2011).

The selectivity of sensory representations can be modulated dynamically by changes in behavioral state, allowing an animal to learn and respond according to perceptual task. In mammals, the noradrenergic system originating in the locus coeruleus (LC) is implicated in signaling behavioral states such as attention, arousal and expectation (Sara and Bouret 2012; Aston-Jones and Cohen 2005).

In insects, octopamine (OA), structurally and functionally similar to noradrenalin (NA) in mammals (Roeder, 2005), can mediate changes in behavioral state that often promote activity, for example: sensitization of reflex actions in locusts (Sombati and Hoyle 1984); aggressive state in crickets (Stevenson et al. 2005); initiation and maintenance of flight state (Brembs et al. 2007; Suver et al. 2012); and enhanced excitability of *Drosophila* motion detection neurons during flight (Strother et al. 2018). Another role of OA is as a reward signal: a single OA neuron, VUMmx1, mediates the reinforcing function of unconditioned stimulus in the honeybee proboscis extension reflex (Hammer and Menzel 1998; Hammer 1993; Menzel 2012). In *Drosophila*, acquisition of appetitive memory is impaired in *TβH* mutants, unable to synthesize OA (Schwaerzel et al. 2003), and activation of OA neurons can substitute reinforcing stimulus in appetitive learning. Moreover, OA receptors are necessary for reward learning in *Drosophila* (Burke et al. 2012) and crickets (Matsumoto et al. 2015).

To understand the neural mechanisms of OA in higher order sensory discrimination, we used the simple sensory “cortex” of larval *Drosophila*, the calyx, which is the sensory input region of the mushroom bodies (MBs), the insect memory center. Here, each MB neuron (Kenyon cell, KC) typically arborizes in several glomeruli, most of which are organized around the terminus of an olfactory projection neuron (PN); KCs thus combinatorially integrate multiple sensory input channels (Masuda-Nakagawa et al. 2005) and are coincidence detectors of multiple inputs. The APL provides inhibitory feedback (Lin et al. 2014; Masuda-Nakagawa et al. 2014) and helps to maintain KC sparse responses and odor selectivity (Honegger et al. 2011), analogous to inhibition in the mammalian PCx (Poo and Isaacson 2009; Stettler and Axel 2009; Gire et al. 2013). Thus, odors are represented as a sparse ensemble of KCs that are highly odor selective, a property beneficial for memory (Olshausen and Field 2004).

In addition, the larval MB calyx is innervated by two OA neurons, sVUMmd1 and sVUMmx1, ventral unpaired medial neurons with dendritic fields originating in the mandibular and maxillary neuromeres, respectively, of the SEZ in the 3^rd^ instar larva (Selcho et al. 2014). sVUMmd1 and sVUMmx1 are named as OANa-1 and OANa-2, respectively, in the EM connectomic analysis of a 6-hour first instar larva (Eichler et al. 2017; Supp. Fig. 3 of Saumweber et al. 2018). These sVUM1 neurons also innervate the first olfactory neuropile of the antennal lobe (AL). This pattern of innervation is conserved in other insects, for example, the dorsal unpaired median (DUM) neurons in locusts (Reviewed by Bräunig and Pflüger 2001), the VUMmx1 neuron in honeybees (Hammer 1993; Schröter et al. 2007), and OA-VUMa2 neurons in adult *Drosophila* (Busch et al. 2009). In adult *Drosophila*, OA-VUMa2 neurons show also a dense innervation of the lateral horn, implicated in innate behaviors (Busch et al. 2009). The widespread innervation of the insect olfactory neuropiles also resembles the widespread NA innervation of mammalian olfactory processing areas, such as the olfactory bulb, and piriform cortex, by LC neurons originating in the brainstem.

We characterized the innervation pattern and synaptic targets of sVUM1 neurons in the calyx, with MB intrinsic and also extrinsic neurons, the localization of the OA receptor Oamb in the calyx circuit, and the impact of sVUM1 neuron activation on behavioral odor discrimination. For this we used an appetitive conditioning paradigm, and tested the ability of larvae to discriminate between similar odors, as opposed to dissimilar odors. Since the larval connectome is based on a single brain, at first instar stage before octopaminergic connections have become as extensive as at third instar, and to obtain a comprehensive understanding of the synaptic targets of sVUM1s in the third-instar larval calyx, we extended our analysis to previously unanalyzed connectivity of VUM1s, to APL and PNs. Further, we combined light microscopy of third-instar larvae with the connectome described by (Eichler et al. 2017).

We find that sVUM1 neurons in third-instar larvae contact all the major classes of calyx neuron to some degree, consistent with EM synaptic analysis of the 6-hour larva (Eichler et al. 2017). A GFP fusion of the OA receptor OAMB is localized in the terminals of PNs, and activating a subset of 5 SEZ neurons, including sVUM1 neurons, can affect discrimination of similar odors, without affecting underlying olfactory learning and memory ability. We suggest a broad modulatory effect of sVUM1 neurons in the calyx, including a potential role in modulating PN input at the second synapse in the olfactory pathway.

## Results

### sVUM1 neurons in the third-instar calyx and their polarity

#### Two OA neurons innervate throughout the calyx without obvious regional preference

At the third-instar stage, the larval calyx is innervated by two classes of OA neurons, sVUMmd1, and sVUMmx1, originating from the mandibular and maxillary neuromeres, respectively, in the subesophageal zone (SEZ), and labeled by the *Tdc2-GAL4* line (Selcho et al. 2014). These neurons are named OANa-1 and OANa-2, respectively, in the 6-hour larval brain EM connectomic analysis, described by Eichler et al. (2017). OAN-a1 corresponds to VUMmd1, and OAN-a2 to VUMmx1, judging from the respective anterior and posterior positions of their cell bodies in Extended Fig 5 of Eichler et al. (2017). Both neurons are labeled by *GMR34A11-GAL4* (Supplementary Fig. 3 of Saumweber et al. 2018), which we used below in our behavior analysis. To visualize the innervation pattern of these neurons together in the more mature third-instar larva, we used the Multicolor FlpOut technique (Nern et al. 2015). Flies of genotype *pBPhsFlp2::PEST(attP3); HA_V5_FLAG_OLLAS* were crossed to flies of *Tdc2-Gal4*, and single cell clones were generated by heat shock. Each sVUM1 neuron ramified throughout the calyx, and we only ever found a single sVUMmd1 or sVUMmx1 neuron labeled. When both sVUM1 neurons were labeled, they ramified through the calyx in a non-overlapping pattern; no fasciculation between the processes of the two neurons was observed, and each innervated the whole calyx without obvious regional preference, as shown in the 3D image (Fig. 1A). Two cell bodies were labeled in the same channel in the mandibular neuromere, but only one innervated the calyx. A single sVUM1 neuron was identifiable by labeling in a single channel in the maxillary neuromere (Fig. 1B).

**Figure 1.**
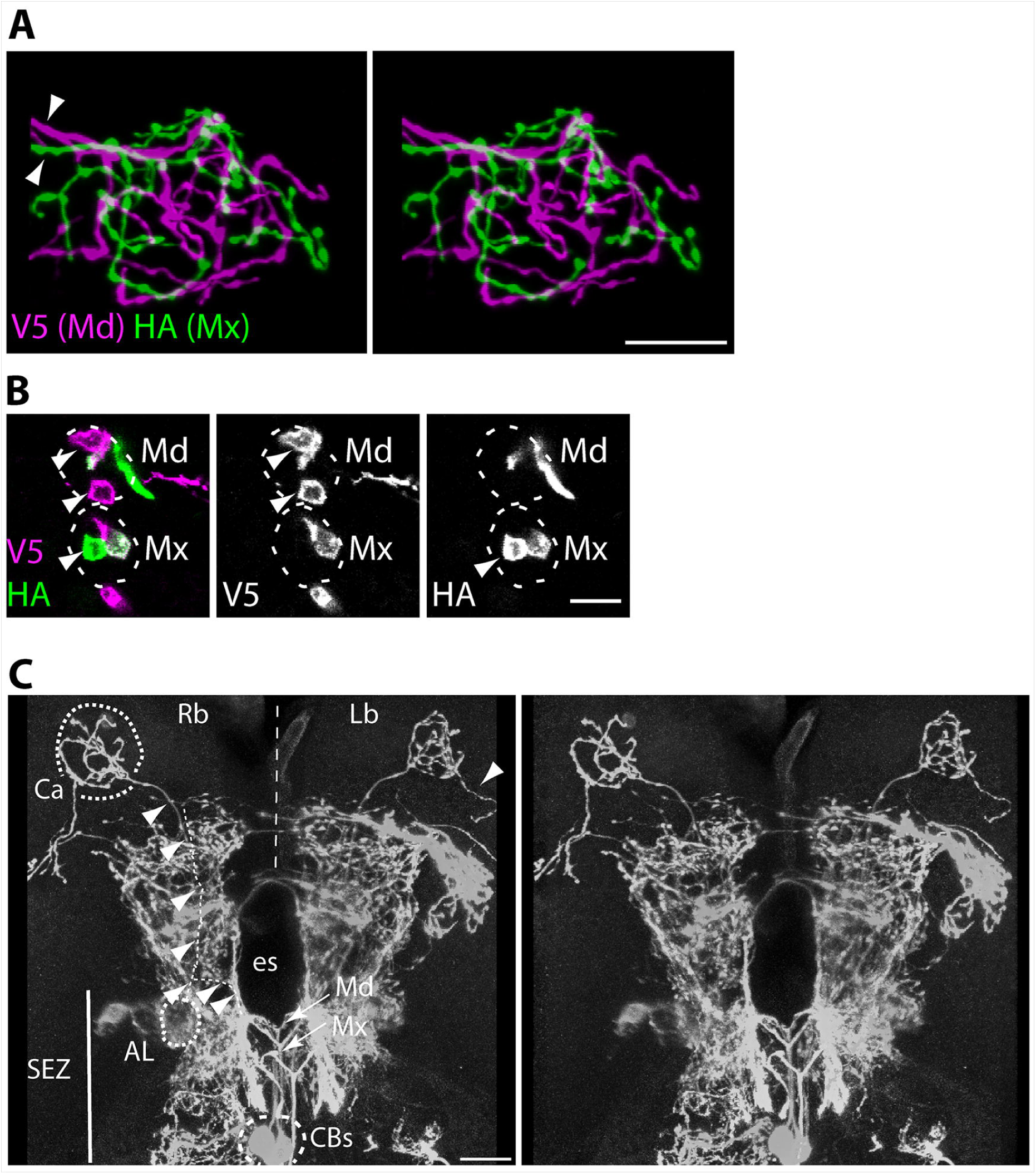
MultiColor FlpOut of *Tdc2-GAL4*-expressing neurons labels two calyx-innervating neurons in third-instar larvae. A-C. Clones of larvae of genotype *w, pBPhsFlp2::PEST; Tdc2-GAL4; UAS-Ollas (attP2) UAS-HA-UAS-V5-UAS-FLAG (attP1)*, were generated by heat shock induction of FLP, and identified by multicolor labeling. **A**. 3D stereo pair of images of an sVUM1Md neuron (V5 tag) and an sVUM1Mx neuron (green, HA tag), both ramifying throughout the calyx. Arrowheads show the axons of the two neurons. Scale bar, 10 µm. **B**. Cell bodies in the SEZ of the same larva. In the mandibular (top) neuromere, two cell bodies labeled in magenta can be identified labeled by antibody to V5 (arrowheads). One of these is the sVUM1md neuron. In the maxillary (bottom) neuromere, a single cell body is only green, identified by anti-HA (arrowhead). **C**. 3D stereo pair of images of a frontal view of a larva in which neurons are clonally labeled with anti-V5, for anatomical clarification. An sVUM1 neuron trajectory is shown. Cell bodies (CBs) of sVUM1 neurons are at the ventral midline of the subesophageal zone (SEZ). Each sVUM1 neuron sends a primary process that bifurcates into secondary processes at the level of the mandibular (Md, arrow) for sVUM1Md, and maxillary (Mx, arrow) neuromere for sVUM1Mx. Each secondary process joins an ascending tract on each side of the esophagus foramen (es), to innervate the ipsilateral protocerebrum, right brain (Rb) or left brain (Lb). While one branch innervates the AL (antennal lobe), another branch separates to follow a tract to the calyx (Ca). Ca and Al are indicated by dotted lines. The AL can be appreciated at the ventral-anterior region of the brain in the 3D. An arrowhead shows a ventral protrusion from the calyx. The trajectories of the VUM neurons are labeled by arrowheads and smaller dotted lines. Scale bar, A, B, 15 µm; C, 20 µm.

Cell bodies of sVUM1 neurons are at the midline of the CNS in the SEZ; their primary processes project dorsally and bifurcate just before the esophagus foramen, to generate two laterally oriented secondary processes each of which joins an ascending tract to the protocerebrum on each brain hemisphere. At the posterior of the AL, the process generates a branch that goes anteriorly and ramifies in the AL. The main branch ascends through the inner antennocerebral tract to reach the calyx at the dorsal protocerebrum. One branch emanates from the calyx and projects ventrally, presumably to the lateral horn (Fig. 1C).

#### OA neurons are presynaptic in the calyx

To visualize the polarity of the calyx-innervating sVUM1 neurons, we used *Tdc2-GAL4* to express either plasma membrane, presynaptic or dendritic markers (Fig. 2). The projections of both sVUMmd1 and sVUMmx1, when visualized by the plasma membrane marker CD4::tdTom, showed dense ramification throughout the calyx, with discrete and abundant bouton-like enlargements along the axonal process, predominantly among glomeruli and in the core of the calyx (Fig. 2A). The presynaptic nature of the *Tdc2-GAL4*-expressing calyx boutons was further supported by the localization of the presynaptic markers nSyb::GFP (Fig. 2A) and Syt::GFP (Fig. 2B), prominently between glomeruli or in the non-glomerular core of the calyx. Only a few dots are visible of the dendritic marker DenMark (Fig. 2B) in the terminals throughout the calyx, On the other hand, DenMark::mCherry is strongly localized in the SEZ region (Fig. 2C), where the postsynaptic processes of VUM neurons are localized. The DenMark::mCherry labeling includes dense arborizations of other *Tdc2-GAL4*-expressing neurons as well as of the sVUM1 neurons. The innervation of the calyx by *Tdc2* neurons can be visualized in Fig. 2D.

**Figure 2.**
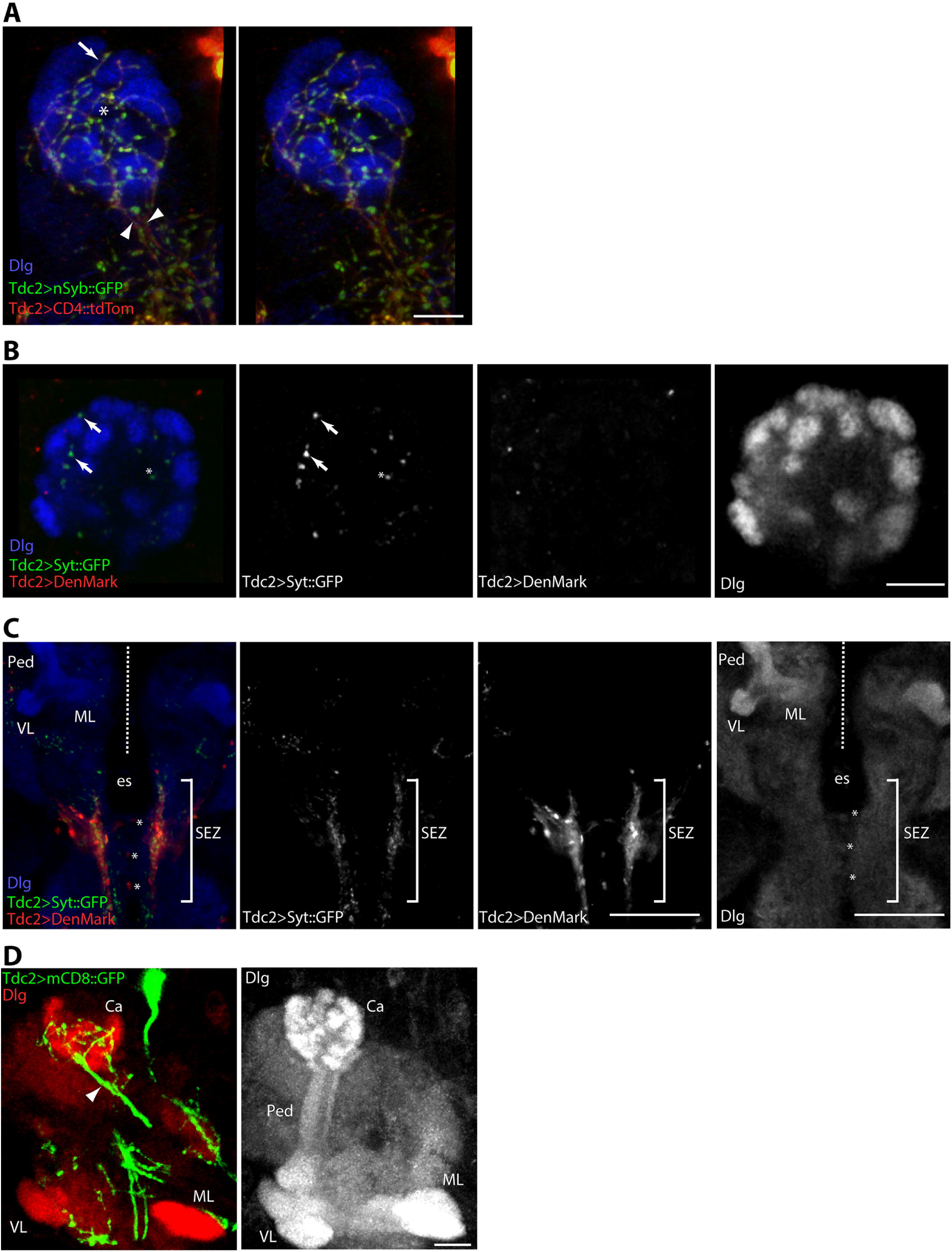
Polarity of calyx innervation by *Tdc2-GAL4-*expressing neurons in third-instar larvae. **A**. 3D stereo pair of images of a third-instar calyx expressing nSyb::GFP and CD4::tdTom using *Tdc2-GAL4*, obtained from a cross between *UAS-CD4::tdTomato (II); UAS-nSyb::GFP (III)* and *Tdc2-GAL4* parents. Two axonal tracts (arrowheads) enter the calyx and ramify throughout it (CD4::tdTom), while processes show nSyb::GFP in boutons (green). Note boutons at the borders of glomeruli (e.g. arrow; glomeruli labeled with anti-Dlg) and core of the calyx (e.g. asterisk). **B**. Confocal section of a larval calyx expressing DenMark::mCherry and Syt::GFP using *Tdc2-GAL4*, obtained from a cross of genotype *Tdc2-GAL4* to BDSC stock 33065. Note the almost complete absence of DenMark::mCherry within the calyx. Asterisk indicates the core of the calyx with adjacent Syt::GFP puncta. Arrows indicates Syt::GFP boutons between glomeruli. **C**. Confocal section of a larval brain expressing Syt::GFP and DenMark::mCherry using *Tdc2-GAL4*. The primary processes of *Tdc2-GAL4-*expressing neurons in the SEZ are shown (asterisks). Brain neuropiles are labeled by Dlg. Dotted line, midline of the CNS between right and left brain hemispheres; es, esophagus; VL, MB vertical lobe; ML, MB medial lobe; SEZ, subesophageal zone (brackets). **D**. Frontal view of a right brain hemisphere, showing an MB calyx innervated by *Tdc2-GAL4* driving expression of mCD8::GFP. Left: projection of 6 sections showing the *Tdc2-GAL4* tract entering the calyx (Ca), arrowhead. Right: the same projection of the MBs labeled by Dlg, right brain hemisphere. Notice the glomerular structure of the calyx. **A, B**: Anterior to the bottom, medial to the right, right brain orientation. Scale bars, 10 µm. **C**: Ventral view, anterior to the top. Scale bar, 50 µm. **D**: Frontal view, anterior to the bottom, medial to the right. Right brain. Scale bar, 10 µm.

**Figure 3.**
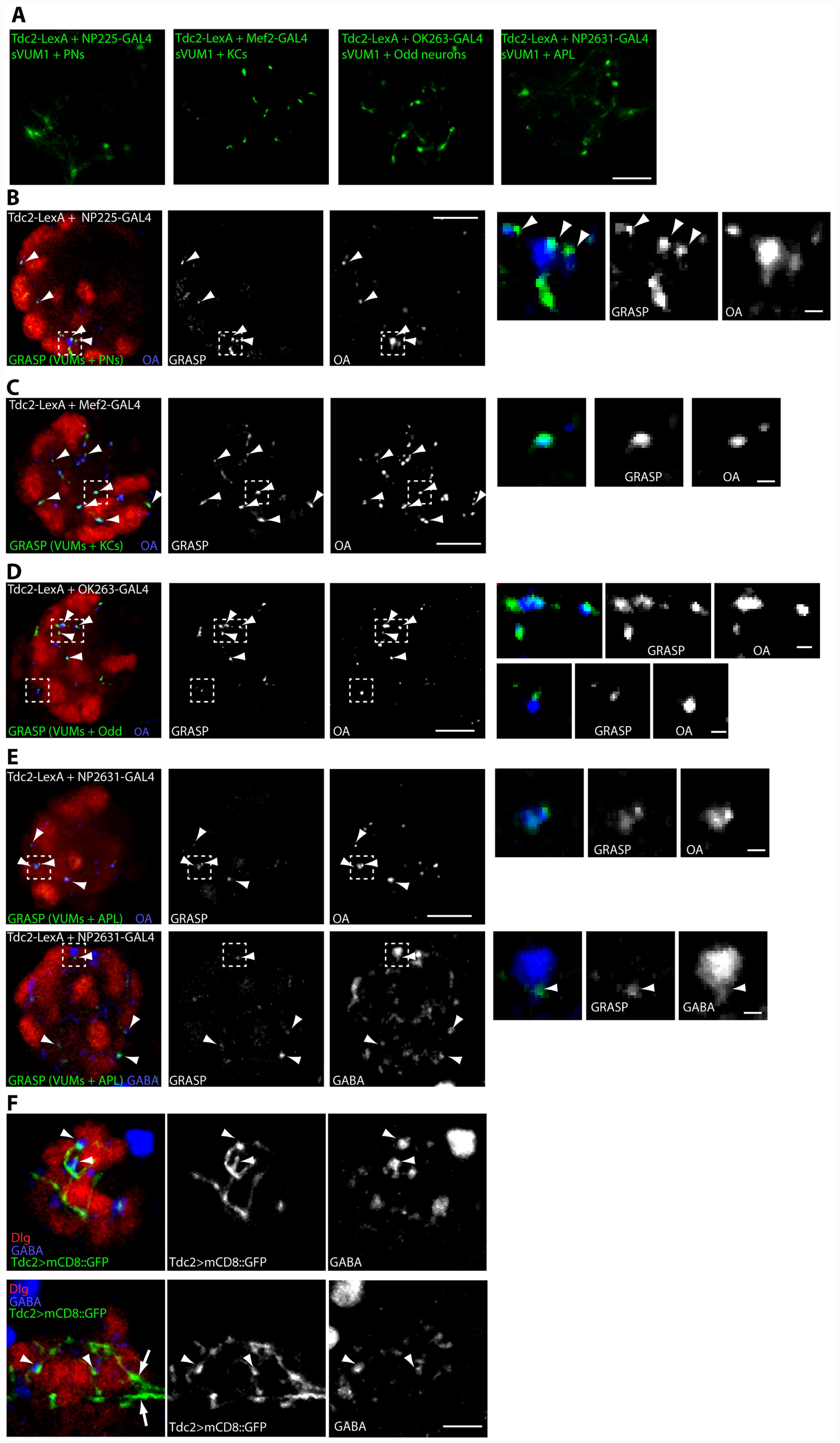
GRASP shows calyx contacts of sVUM1 termini with PNs, larval APL, and *odd-*like neurons, and few KCs. A line carrying GRASP constructs *UAS-CD4::spGFP1-10* and *LexAop-CD4::spGFP11*, was crossed to each of the following lines as needed: *NP225-GAL4(II); Tdc2-LexA(III)*, or *OK263-GAL4(II); Tdc2-LexA(III)*, or *NP2631-GAL4(II); Tdc2-LexA(III)*, or *Tdc2-LexA(II); Mef2-GAL4(III), and GRASP signals were detected in the larval progeny* **A**. GRASP signal between sVUM1 neurons (expressing *Tdc2-LexA*) and other calyx neurons expressing the *GAL4* lines shown, is detected as native GFP fluorescence in confocal sections. Scale bar, 10 µm. **B-E**. GRASP signal between sVUM1 neurons (expressing Tdc2-LexA) and other calyx neurons expressing the *GAL4* lines shown, detected by rat monoclonal anti-GFP. Examples of GRASP signal are highlighted with arrowheads; areas inside broken square are shown at higher zoom. sVUM1 termini are labeled using anti-OA, calyx glomeruli using anti-Dlg. Scale bars in main panels 10 µm, in insets 1 µm, here and throughout figure. **B**. GRASP signal between sVUM1 termini and PNs, localized mainly around glomeruli. OA signals without GRASP are localized to the calyx core. Examples of GRASP signal are highlighted with arrowheads. GRASP signal overlaps partially with sVUM1 termini labeled with OA (inset). Areas inside broken lines, here and subsequently, are shown at higher zoom. **C-E**. GRASP signals between sVUM1 termini and KCs (**C**), Odd neurons (**D**), or the APL neuron (**E**). **E**. GRASP signals between sVUM1 and APL neurons are localized around glomeruli, or in the core of the calyx. Upper panels show the extent of GRASP overlap with OA-containing sVUM1 terminals. Lower panels show some GRASP signals overlapping partially with large GABA boutons. **F**. Confocal sections of a third-instar calyx expressing *mCD8::GFP* using *Tdc2-Gal4*, labeled with anti-GABA and anti-Dlg. Top is a dorsal section of calyx, bottom is a ventral section of the same calyx. Arrowheads indicate GABA boutons apposed to or overlapping with *Tdc2* neuron terminals. Arrows indicate the two axonal tracts of *Tdc2* neurons innervating the calyx.

### Identifying third-instar calyx OA neuron partners

To obtain a comprehensive synaptic connectivity between sVUM1 neurons and intrinsic as well as all extrinsic neurons innervating the calyx, we used *Tdc2-LexA*, along with *GAL4* lines expressing in other calyx neurons, to drive the expression of the GFP Reconstitution Across Synaptic Partners (GRASP, Gordon and Scott 2009) constructs *LexAop-CD4::spGFP11 and UAS-CD4::spGFP1-10* (Fig. 3). We labeled olfactory PNs using *NP225-GAL4* (Tanaka et al. 2004), and KCs using *Mef2-GAL4* (Zars et al. 2000). We also tested for GRASP between sVUM1 neurons and two other classes of extrinsic calyx neurons. First, we labeled the larval APL using *NP2361-GAL4* (Masuda-Nakagawa et al. 2014). Second, two of the “Odd” class of neurons that arborize throughout the calyx (Slater et al. 2015) have been designated MBON-a1 and MBON-a2 by Eichler et al (2017); we identified a *GAL4* line, *OK263-GAL4*, which expresses in these neurons.

#### GRASP fluorescence

We detected GRASP using GFP fluorescence as widely distributed puncta in the calyx, in live images of brains, suggestive of specific synaptic connections, between the sVUM1 neurons on the one hand, and PNs, KCs, the Odd, and APL neurons on the other (Fig. 3A). Control crosses expressing the GRASP constructs under control of either a *GAL4* or *LexA* calyx driver alone showed almost no GRASP puncta (Supplementary Fig. 1), showing the specificity of our rat monoclonal anti-GFP for reconstituted GFP. These findings suggest that at the third-instar larva, the sVUM1 neurons may form synapses with all the neuronal classes that innervate throughout the calyx: PNs, KCs, the APL, and Odd neurons.

To test whether GRASP signals represented synaptic contacts of the sVUM1 neurons, we also immunolabeled brains with anti-OA. GRASP signals were identified using a criterion that each GFP signal was observed in at least two consecutive confocal sections, and discounting occasional GFP-positive axonal tracts that were negative for OA. As noted below, the large majority of GFP puncta either overlapped or were directly apposed to OA signals, suggesting that GFP puncta were mostly or entirely specific for synaptic contacts, and did not form widely at non-synaptic contacts.

#### OA termini synapse with PNs

Using *Tdc2-LexA* and *NP225-GAL4* to express the GRASP constructs, we found GRASP signal at 49 ± 3% of 80 ± 4 (n=5) OA-positive boutons. OA-positive GRASP signals (Fig. 3B) were found in the core of the calyx away from glomeruli, in interglomerular spaces, and along the periphery of glomeruli. They are likely contacts between sVUM1 termini and PN axons. Almost all GFP puncta (87 ± 2%, n=5) overlapped with or were apposed to OA boutons and are therefore potential synaptic contacts of the sVUM1 neurons; the remaining 13% could represent non-synaptic contacts, or synapses of PNs onto postsynaptic sites on the sVUM1 neurons. Therefore, PNs are commonly postsynaptic to the termini of sVUM1 neurons, on their axonal or presynaptic processes, making axon-axon synapses.

#### OA termini synapse with KCs in the calyx

Using *Tdc2-LexA* and *MB247-GAL4* to express the GRASP constructs, we found GRASP signals at 51 ± 4 % of 185 ± 34 (n=3) OA boutons in each calyx (Fig. 3C). GRASP puncta overlapping with OA were found in the interglomerular space. Again, most GFP puncta (86 ± 1%) overlapped with or were apposed to OA. Since third-instar larvae are estimated to have 250-300 KCs per brain hemisphere (Pauls et al., 2010), our data suggest that sVUM1 neurons together synapse onto up to about one third of third-instar KCs.

#### OA termini synapse with Odd neurons in the calyx

Using *Tdc2-LexA* and *OK263-GAL4* to express the GRASP constructs, we found GRASP signals at 51 ± 0.3 % of 124 ± 4 (n=3) OA labeled boutons (Fig. 3D). Again, the majority of GFP puncta (80 ± 1%) overlapped with or were apposed to OA, suggesting that Odd neurons are mostly postsynaptic to sVUM1s.

#### OA termini synapse with the APL

Using *Tdc2-LexA* and *NP2361-GAL4* to express the GRASP constructs, we found GRASP signal at 77 ± 4% of 74 ± 12 (n=4) OA terminals, indicating that sVUM1 neurons are presynaptic to the larval APL (Fig. 3E top panels). Most of these GRASP signals were found between glomeruli, and more abundant towards the ventral calyx.

Around 67 ± 3% of GFP (n=3) signals overlapped with OA, and therefore likely represent synapses of the sVUM1 neurons onto the APL. A higher frequency of GFP puncta did not overlap with OA (33 ± 3%) than was the case for other calyx neurons; this could potentially be due to synapses of the APL onto OA neuron axons. In support of this, we found some GABA termini in close proximity to GRASP (Fig. 3E, bottom panels); labeling of Tdc2-GAL4>mCD8::GFP calyces with anti-GABA also showed some apposition of sVUM1 boutons to GABAergic termini of the APL (Fig. 3F).

#### Single cell GRASP

While the above GRASP experiments reveal the partners of the sVUM1 neurons in the calyx, they do not reveal whether the sVUMmd1 and sVUMmx1 have different partners. We therefore performed single-cell GRASP to label the contacts of each sVUM1 randomly, using Brp::mCherry as a presynaptic marker to verify whether GRASP signals have a synaptic localization. We distinguished the two sVUM1 neurons by the positions of their cell bodies in the SEZ, using local neuropil landmarks revealed by anti-Dlg labelling (Fig. 4A), and using single cell clones, we could identify sVUMmd1 and sVUMmx1 individually (Fig. 4B).

**Figure 4.**
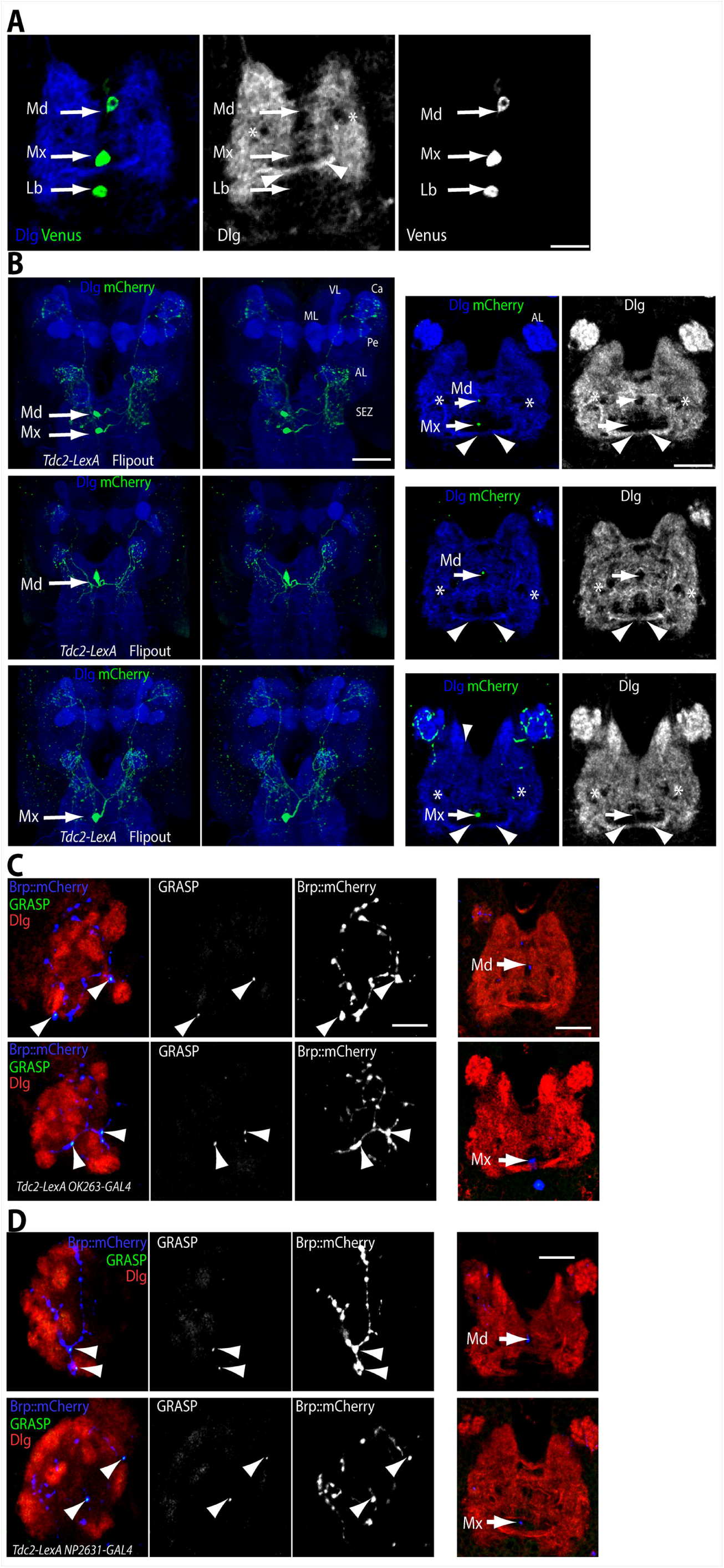
Single cell GRASP of sVUM1 neurons in the calyx. **A**. Identification of sVUM1 neurons based on SEZ anatomical landmarks. Males of genotype *w/Y*; *Tdc2-LexA(II)*; *GMR34A11-GAL4(III) / TM6B* were crossed to females of genotype *Tub84B(FRT-GAL80)1, w*; *LexAop2-FLPL(II)*; *UAS-Chrimson*.*mVenus(III) / TM6B*. The larval progeny of this cross, of genotype *FRT*.*GAL80 w / (w or Y)*; *Tdc2-LexA(II) / LexAop2-FLPL(II)*; *GMR34A11-GAL4(III)* / *UAS-CsChrimson*.*mVenus(III)*, express Venus in cells that express both *Tdc2-LexA* and *GMR34A11-GAL4*. sVUM1md neurons (Md) are in a small anterior medial Dlg-negative gap, with bilaterally symmetrical gaps (asterisks) nearby. sVUM1mx neurons (Mx) are localized in a large medial gap that lies just anterior to a commissural DLG pathway (arrowheads). sVUMlb neurons (Lb) are localized posteriorly to this commissural pathway. Scale bar, 24 µm. **B-D**. Single-cell GRASP was induced between sVUM1 neurons expressing *Tdc2-LexA*, and potential calyx partner neurons that expressed different *GAL4* lines. sVUM1 neurons were individually labeled by heat-shock-induced FlipOut of larvae of genotype *P{hsFLP}12/(w or Y); GAL4(II) / +; Tdc2-LexA(III) / LexAOp2-IVS>stop>spGFP11::CD4::HA-T2A-Brp::mCherry (attP2), UAS-spGFP1-10::CD4, UAS-HRP::CD2*. **B**. Single cell clones, expressing Brp::mCherry in subsets of *Tdc2-LexA*-expressing neurons; each row shows a separate clone. Left image pairs are stereo views of reconstructions showing the entire trajectory of labelled neurons; MB calyx (Ca), pedunculus (Pe), and medial and vertical lobes (ML, VL), and the antennal lobe (AL) and subesophageal zone (SEZ) are labeled in the top row. Right images are single confocal sections through the SEZ of the same larvae, showing the primary process of each labelled sVUM1 (arrows) and the anatomical landmarks shown in **A**. First row, sVUMmd1 and sVUMmx1. Second row, sVUMmd1. Third row, sVUMmx1. Scale bar is 45 µm in 3D images (Left), 30 µm in SEZ sections (Right). **C-D**. Single-cell GRASP between sVUM1 neurons expressing *Tdc2-LexA*, and (**C**) Odd neurons expressing *OK263-GAL4*, or (**D**) larval APL expressing *NP2631-GAL4*. Left panels: larval calyx labelled with anti-DsRed to visualize Brp::mCherry (enriched at presynaptic sites but not confined to them), anti-Dlg to label neuropile, and anti-GFP to visualize GRASP signals. Arrowheads indicate GRASP signals. Right panel: region of same brain labelled with anti-Dlg to visualize neuropile, and anti-DsRed to visualize the VUM neurons. Top rows show sVUMmd1 (Md), bottom rows show sVUMmx1 (Mx). Arrows indicate the primary process of each sVUM1 neuron. Scale bars: 10 µm.

**Figure 5.**
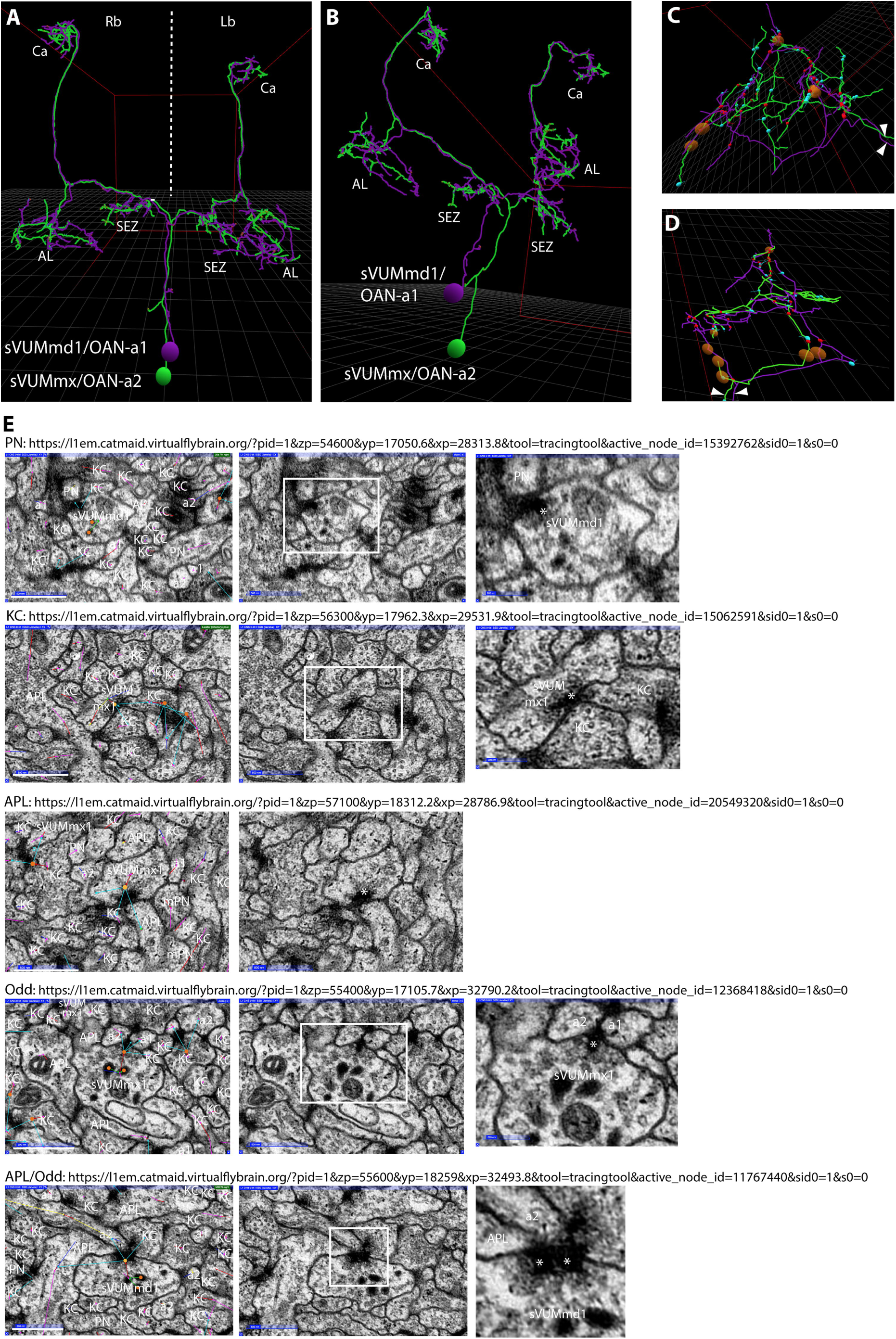
Synapses of sVUM1 neurons onto other first-instar calyx neurons. **A**. Ventral view of a reconstruction of sVUMmd1/OAN-a1 (magenta) and sVUMmx1/OAN-a2 (green), including their dendritic arborizations in the subesophageal zone (SEZ) and presynaptic terminals in the antennal lobe (AL) and calyx (Ca). Right brain (Rb) is to the left of the midline (dotted line), and left brain (Lb) to the right. sVUM1 neurons bifurcate at the midline and a single neuron innervates both brain hemispheres. **B**. A ventrolateral view of the same reconstructions as A. **C**,**D**. Reconstructions of sVUM1md1 and sVUMmx1 calyx projections in the right (**C**) and left (**D**) brain. Red circles are sVUM1 presynaptic termini (blue circles are postsynaptic sites). Note the 39 presynaptic terminals of sVUM1 neurons in the right calyx (C) and the 28 presynaptic termini in the left calyx (D). Brown circles are tracing sites not finished. Images were generated by analysis of neuron tracing using the 3D tool of CATMAID on the publicly available first-instar larval connectome on the Virtual Fly Brain site (https://l1em.catmaid.virtualflybrain.org; Licence CC-BY-SA_4.0). **E**. EM sections of first-instar calyx (Eichler et al., 2017) showing synaptic partners of either sVUMmd1 or sVUMmx1, named as OAN-a1 and OAN-a2 by Eichler et al. (2018) and in CATMAID as “Anterior Ladder” and “Posterior Ladder” neurons, respectively. Sections were visualized using CATMAID software via the Virtual Fly Brain site. Each row shows a different example of postsynaptic target neurons: a PN; a KC; the APL (labeled in CATMAID as MBE12); Odd neurons (MBON-a1 and MBON-a2, labeled here as a1 and a2, and in CATMAID as MBE7a and MBE7b); and a tripartite synapse of an sVUM1 neuron on both an Odd neuron (MBON-a2) and the APL. In each row, the first (left) panel shows a section with CATMAID annotation, and the identities of neurons close to the sVUM1 neuron and its synaptic target. Notice the connector (orange filled circle) placed on the sVUM1 neuron and the linked cyan arrows projecting to postsynaptic partners. Red arrows pointing to the connector indicate that the connector is on the presynaptic neuron. The second panel shows the same image with annotation omitted to allow better visualization. The third panel (where present) shows a magnified view of the inset in the middle panel; a presynaptic release site in the sVUM1 neuron, characterized typically by a T-bar and surrounding synaptic vesicles, is shown with an asterisk. Scale bars: white lines, 500 nm. URLs link to the locations of images shown.

GRASP signals were detected between the *odd*-expressing neurons and both sVUMmd1 and sVUMmx1 (Fig. 4C), and similarly between the larval APL, and both sVUMmd1 and sVUMmx1 (Fig. 4D). Compared to standard GRASP, single cell GRASP signals were fewer, but clearly present and overlapping with the presynaptic marker Brp::mCherry (Fig. 4D). The main targets of VUMmd1 and VUMmx1 in the olfactory pathway are the antennal lobe (AL), and calyx, as described in Selcho et al. (2014). There is also a prominent branch innervating the basolateral protocerebrum (anteromedial to the AL) around the ventral midline at the esophagus foramen (as defined for adult flies, Busch et al. 2009) in the brain. Therefore, at least as judged by the APL and Odd neurons, both sVUM1 neurons appeared to have similar targets in the calyx.

#### Comparison with first instar larva calyx synapses

The enlargements seen using synaptic markers in Fig. 2A contained OA (Fig. 3), therefore they are presynaptic boutons. Using the online publicly available first instar connectome (https://l1em.catmaid.virtualflybrain.org) on the Virtual Fly Brain site (Osumi-Sutherland et al. 2014; Cantarelli et al. 2018), we analyzed the synapse distribution and synaptic partners of the sVUM1 neurons specifically in the first instar larva calyx, not previously analyzed. We generated a tracing representation of the sVUM1 neurons (Fig. 5A,B) using the 3D tool of the publicly available CATMAID software. While the growth, morphology and anatomical organization of the two sVUM1 neurons is similar to the anatomical organization of the third-instar larvae, the arborizations in the calyx are fewer in the first instar compared to third-instar larvae. In our GRASP analysis, we observed 89 ± 7 (mean ± SEM; n= 12) OA-positive boutons per calyx for both sVUM1 neurons, compared to 28 presynaptic synapses marked in the left brain calyx, and around 39 in the right brain calyx in the single 6-hour larva brain connectome (Fig 5C,D).

Moreover, the number of connections between sVUM1 and other neurons was substantially lower at the 1^st^ instar compared to the third-instar stage (Supplementary Table S1), with the first-instar brain having less than 50% of synaptic numbers per calyx per neuron type compared to the third-instar brain.

Given the GRASP signals at sVUM1 synaptic termini, we predicted that we might find the same sVUM1 synaptic targets in the first-instar calyx connectome. Eichler et al. (2017), reported synapses of sVUM1 onto the Odd neurons, and a small fraction of KCs. We found examples of these synapses, and additionally, synapses not yet described between sVUM1 neurons onto PNs and the APL neuron (Fig. 5E). In addition to the apparently simple synapses of sVUM1 neurons onto PNs, KCs, Odd neurons, and the APL, we sometimes found adjacent synaptic contacts of either sVUM1 neuron onto both an Odd neuron and the APL (Fig. 5E), suggesting locally coordinated circuit regulation of both these neurons by the sVUM1 neurons. Clear vesicles and typical synaptic bars were present in sites of connectivity between sVUM1s and other neurons (Fig. 5E), supporting a synaptic release of octopamine. Although sVUM1s also contain dense core vesicles, our studies did not address whether sVUMs release neuropeptides.

### Localization of a genomic Oamb::GFP fusion in PN terminals in calyx

To further understand how and where OA might act in the calyx, we investigated the localization of OA receptors in the calyx. *Drosophila* has a number of OA receptor classes defined both by sequence comparisons and pharmacology. Octopamine receptor in mushroom bodies (Oamb, also known as Dmoa1A or CG3856), an ortholog of human α1-adrenergic receptor (Roeder et al. 2003; Bauknecht and Jékely 2017), is enriched in the MBs (Han et al. 1998). *Drosophila* also has three OctβR receptors, which stimulate cAMP levels (Maqueira et al. 2005; Balfanz et al. 2005).

To detect the expression and subcellular localization of Oamb, we used recombinase-mediated cassette exchange (RMCE) with a MiMIC insertion (Venken et al. 2011), *MI12417*, in the third coding-region intron of *Oamb*, to tag endogenous *Oamb* with an exonic *EGFP-FlAsH-StrepII-TEV-3xFlag* fusion (Supplementary Figs. 2-5). Insertion of an EGFP-encoding exon here should tag all known splice variants of the Oamb protein in their third cytoplasmic loop, downstream of transmembrane (TM) domain 5 (Supplementary Figs. 6,7); this includes the alternative TM6-TM7 regions encoded by two alternative groups of C-terminal exons (Supplementary Figs. 5-7). Therefore, a protein trap generated from the *MI12417* insertion will not disrupt any transmembrane domains.

**Figure 6.**
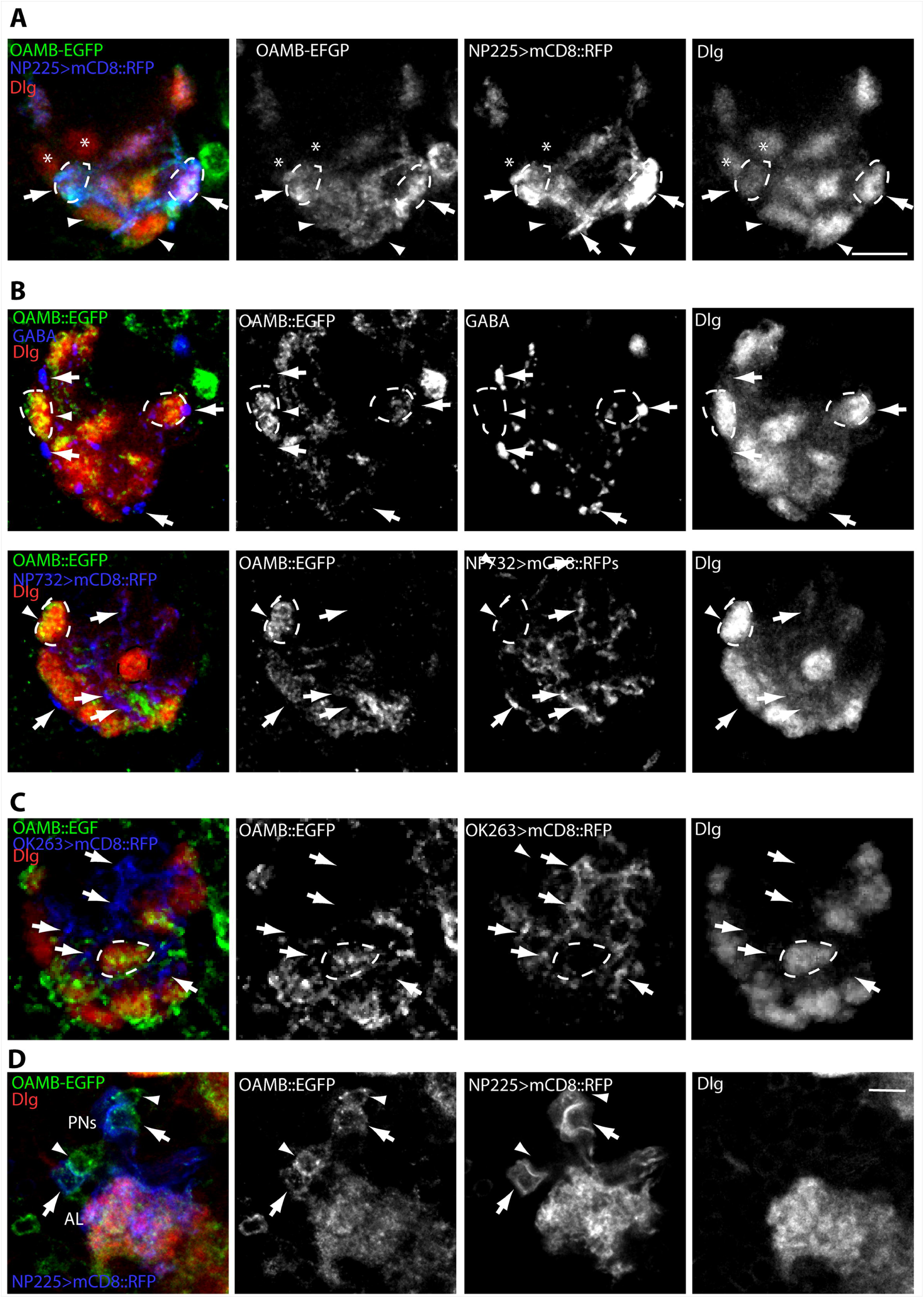
OAMB-EGFP localization in PN presynaptic termini. **A**. Calyces of a larva carrying *Oamb::EFGP, NP225-GAL4* and *UAS-RFP* labeled with chick polyclonal anti-GFP, anti-DsRed and anti-Dlg. Oamb::EGFP localizes to PN terminals in all glomeruli labeled by *NP225-GAL4* (arrows), and to a few glomeruli not labeled by *NP225-GAL4* (arrowheads). A few glomeruli express neither *NP225-GAL4* nor Oamb::EFGP (asterisks). One arrow indicates a PN process. **B**. Top row: Oamb::EGFP does not overlap with APL termini labeled by anti-GABA (example at arrows). Note the prominent GABA boutons without Oamb::GFP at the periphery of glomeruli. Bottom row: APL projections labeled by *NP0732-GAL4*. Varicosities along processes are devoid of Oamb::GFP. **C**. Oamb::EGFP does not overlap substantially with Odd neuron dendrites labeled by *OK263-GAL4*. Prominent varicosities are devoid of GFP (arrows). A glomerulus labeled by Oamb::EGFP does not overlap with Odd neuron dendrites (dotted line). A-C: Representative glomeruli labeled by Oamb::EGFP are indicated by dotted lines and arrows. D. Antennal lobe of a larva carrying *Oamb::EFGP, NP225-GAL4* and *UAS-RFP* labelled with chick polyclonal anti-GFP, anti-DsRed and anti-Dlg. Oamb::EGFP is detected in PN cell bodies (arrowheads), and cell bodies not labelled by *NP225-GAL4* (arrows). Notice the weak diffuse labeling of Oamb::GFP in the AL. Scale bar, 10 µm.

**Figure 7.**
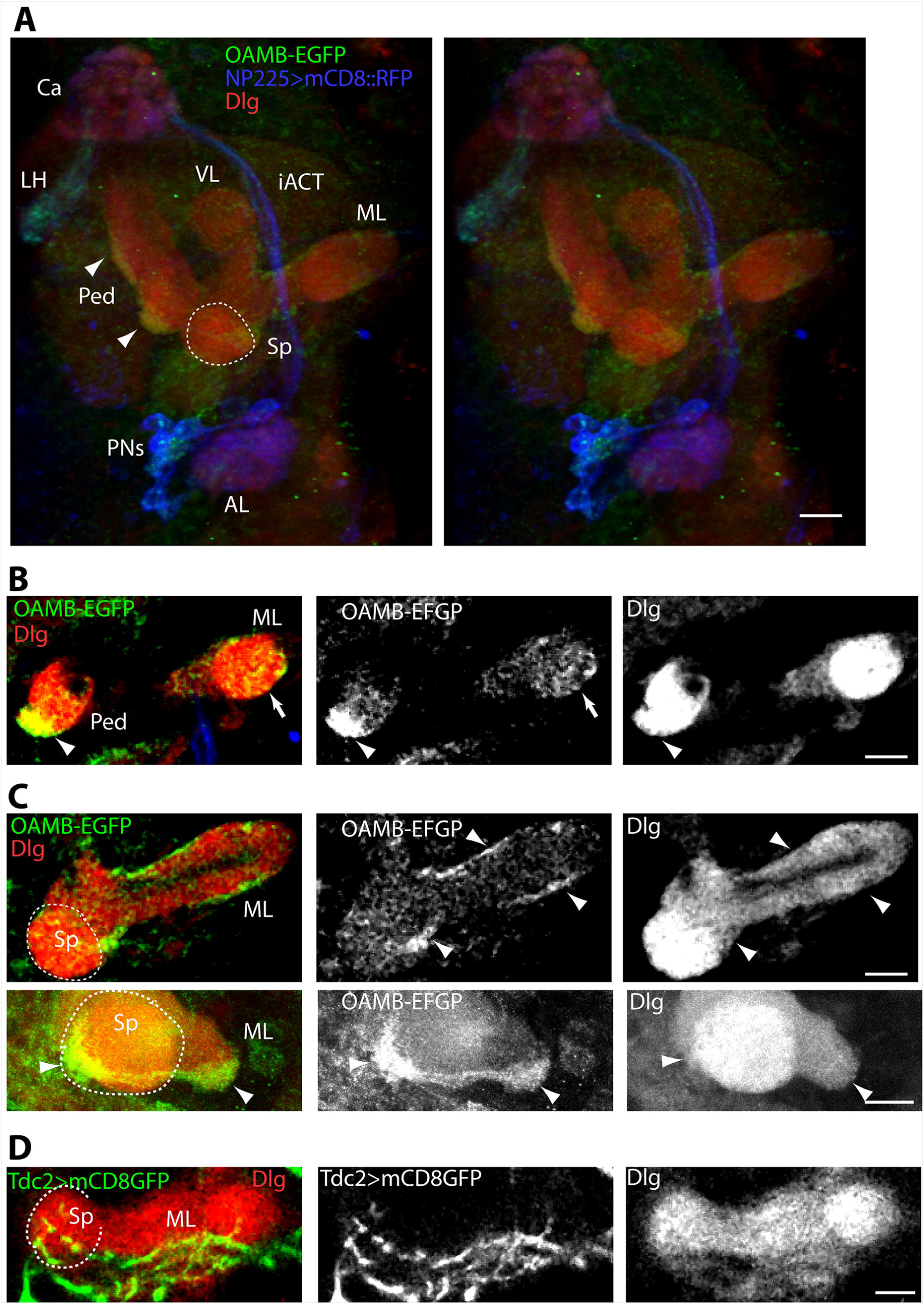
Oamb-EGFP localization in MB pedunculus and lobes. **A**. 3D stereo pair of images of frontal views of a third-instar larval PN pathway in a single brain hemisphere, expressing *Oamb::EFGP, NP225-GAL4* and *UAS-RFP* labeled with chick anti-GFP, anti-DsRed and anti-Dlg. PNs with cell bodies lateral to the AL project to the MB calyx and the LH (lateral horn) via the iACT (inner antennocerebral tract). Oamb::EGFP was detected at the lateral horn (LH) and calyx, lateral pedunculus (arrowheads), and medial lobes (ML). **B**. Confocal section showing Oamb::EGFP localization in lateral pedunculus (arrowhead) and medial lobe (arrow). **C**. Top row: Confocal section showing Oamb::EGFP in the medial lobe (ML) and in the heel of the MB. Bottom row: higher magnification section of Oamb::GFP labeling at the spur (Sp) and medial lobe. **D**. Section through the MB spur and medial lobe in third-instar larva carrying *Tdc2-GAL4* and *UAS-mCD8::GFP*. Notice the GFP-expressing processes at the ventral side of the medial lobe and the spur. The spur region is indicated by dashed lines in A,C,D. Scale bars, 10 µm.

Six recombinant *Oamb::EGFP* stocks were recovered with the EGFP-encoding exon inserted in the same orientation as the *Oamb* transcript (Supplementary Fig. 7). One of these was designated as *Mi{PT-GFSTF*.*1}Oamb*^*MI12417-GFSTF*.*1*^, or *Oamb(MI12417)::EGFP*.*1* or *Oamb::EGFP* for short. Both the original *MI12417 MiMIC* insertion, and *Oamb(MI12417)::EGFP* stocks were homozygous infertile, as expected from the egg-laying defects of *Oamb* mutants (Deady and Sun 2015), suggesting that the Oamb::EGFP fusion might not be a functional Oamb protein. However, Oamb::EGFP was localized to glomeruli in the larval calyx (Fig. 6), implying that the protein folded normally and was not degraded by the ER unfolded protein response. Expression of *UAS-RFP* in the olfactory PN line *NP225-GAL4* showed localization of Oamb::EGFP in all PN termini labeled with the GAL4 line, as well as in some calyx glomeruli not labeled by *NP225-GAL4*, which may be either sites of non-olfactory sensory input, or olfactory glomeruli not labeled by *NP225-GAL4* (Fig. 6A). The restriction of Oamb::EGFP to specific glomeruli implies that it is unlikely to be expressed in KC dendrites in the calyx, which arborize through all glomeruli. We also found no overlap of Oamb::EGFP with GABAergic APL terminals in the calyx (Fig. 6B), implying that it was not expressed in the larval APL. *OK263-GAL4* calyx projections also showed little or no overlap with Oamb::EGFP (Fig. 6C), suggesting that Oamb is not expressed in the Odd neuron calyx dendrites.

In the olfactory pathway Oamb::EGFP was detected diffusely in the AL. Cell bodies of PNs labeled by *NP225-GAL4>mCD8::RFP* also expressed Oamb::EGFP (Fig. 6D), and other neurons surrounding the AL, but not labeled by *NP225-GAL4>mCD8::RFP*, expressed Oamb::EGFP (Fig. 6D). These could be interneurons and potentially the main source of labeling in the AL. No distinct glomeruli were detected. Oamb::EGFP was detected in the lateral horn, along the lateral pedunculus, with strong expression towards the anterior lateral end, where a separate compartment along the pedunculus is labeled (Fig. 7A,B). Oamb::EFGP is detected around the medial lobe and the spur, spur as defined by Younossi-Hartenstein, et al 2003, Fig. 6E, (Fig. 7C). This localization of Oamb::EGFP overlaps with innervation by *Tdc2-GAL4*-expressing neurons in the spur and around the ventral medial lobe (Fig. 7D).

We found no detectable localization of MiMIC GFP-tagged DmOctβR receptors to the calyx (data not shown). Octβ1R::EGFP (*CG6919*) was detected weakly in a few ventral and medial AL glomeruli by a polyclonal anti-GFP, but was not detectable in the calyx. Octβ2R::EGFP (*CG6989*), was not detectable in either the calyx or AL, although it was expressed in a number of adjacent cell bodies that did not colocalize with PNs as labeled by *NP225-GAL4* driving RFP expression. We could not detect Octβ3R::EGFP anywhere in the brain, and therefore the fusion might not be expressed, or misfold and be degraded.

### Activating an OA neuron subset including sVUM1 neurons impairs behavioral odor discrimination

Since the calyx is a site where MB neurons process olfactory information that comprises conditioned stimuli in associative learning, we reasoned that modulating the processing of this information might affect the ability of the brain to discriminate among different conditioned stimuli representations while learning, but without affecting its underlying learning ability. Therefore, to test whether OA innervation of the calyx affected odor discrimination during learning, we developed an assay that could distinguish odor discrimination ability from learning ability (Fig. 8). The rationale of this assay was developed in the honeybee by Stopfer et al. (1997) in pattern recognition. Desynchronization of PN ensembles impaired fine discrimination of molecularly similar odors but not dissimilar odors. When bees are conditioned with one odor and sucrose, the conditioned response generalizes to structurally similar odors used for conditioning (Smith and Menzel 1989). Therefore, we reasoned that mixtures of two odors at different ratios would be more similar and harder to distinguish than the two pure constituent odors, and this would allow us to test changes in the degree of discrimination ability by activation of sVUM1s neurons; this rationale was used successfully by Lin et al. (2014). By combining odor choice with an appetitive learning paradigm (Scherer et al. 2003), we tested the effect on behavioral odor discrimination of optogenetic activation of OA neurons in third-instar larvae, using the long-wavelength absorbing channelrhodopsin, CsChrimson (Klapoetke et al. 2014).

**Figure 8.**
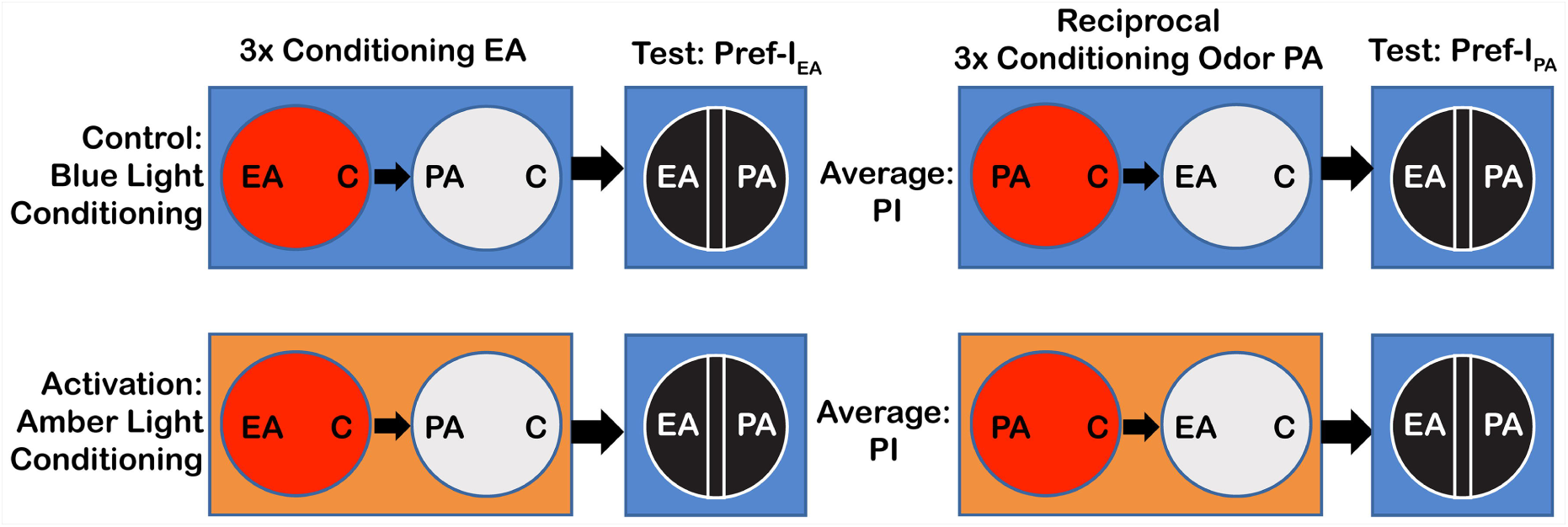
Behavioral discrimination assay and optogenetics. Larvae are conditioned for three cycles of: an agarose fructose plate (red), carrying a container with one odor (here ethyl acetate, EA) and a control container with mineral oil (C), followed by an agarose plate lacking fructose (light gray), but carrying a container with a different odor (here pentyl acetate, PA) and a control container with mineral oil. Larvae are then tested by being placed on an oblong area in the middle of an agarose plate, with a choice of EA or PA, and a black background to provide contrast for counting larvae, and a conditioned preference index for EA (Pref-I_EA_) is measured. A different group of larvae are conditioned and tested with reciprocal odor pairing. Learning is measured as a Performance Index (PI) averaged from Pref-I_EA_ and Pref-I_PA_. Control experiments (top row) are carried out in dim blue light. Optogenetic activation of CsChrimson is accomplished by amber light during the conditioning phases.

Since we did not have *GAL4* or *LexA* drivers completely specific for sVUM1 neurons, we used an intersectional approach to restrict the expression of CsChrimson to a small subset of OA neurons including the sVUM1 neurons. We could use *LexAop-FLP* and a *GAL80* cassette flanked by two *FRT* sites to express *UAS-CsChrimson* only in neurons that expressed both *GMR34A11-GAL4* (which labels some VUM neurons in addition to non-OA neurons), and *Tdc2-LexA*. We thus expressed CsChrimson in only five OA neurons in the SEZ of the larval brain: two in the mandibular neuromere (including sVUMmd1), two in the maxillary neuromere (including sVUMmx1), and one in the labial neuromere (n=8; Fig. 9A). The second neuron labeled in each neuromere could be the sVUM2, characteristic for the lateral branch innervating the optic lobe, the sVUM3 which innervates the dorsal medial protocerebrum and basal medial protocerebrum (Selcho et al. 2014) or an unidentified sVUM; however, sVUM1 neurons are the only neurons of this subset to innervate the AL and calyces.

**Figure 9.**
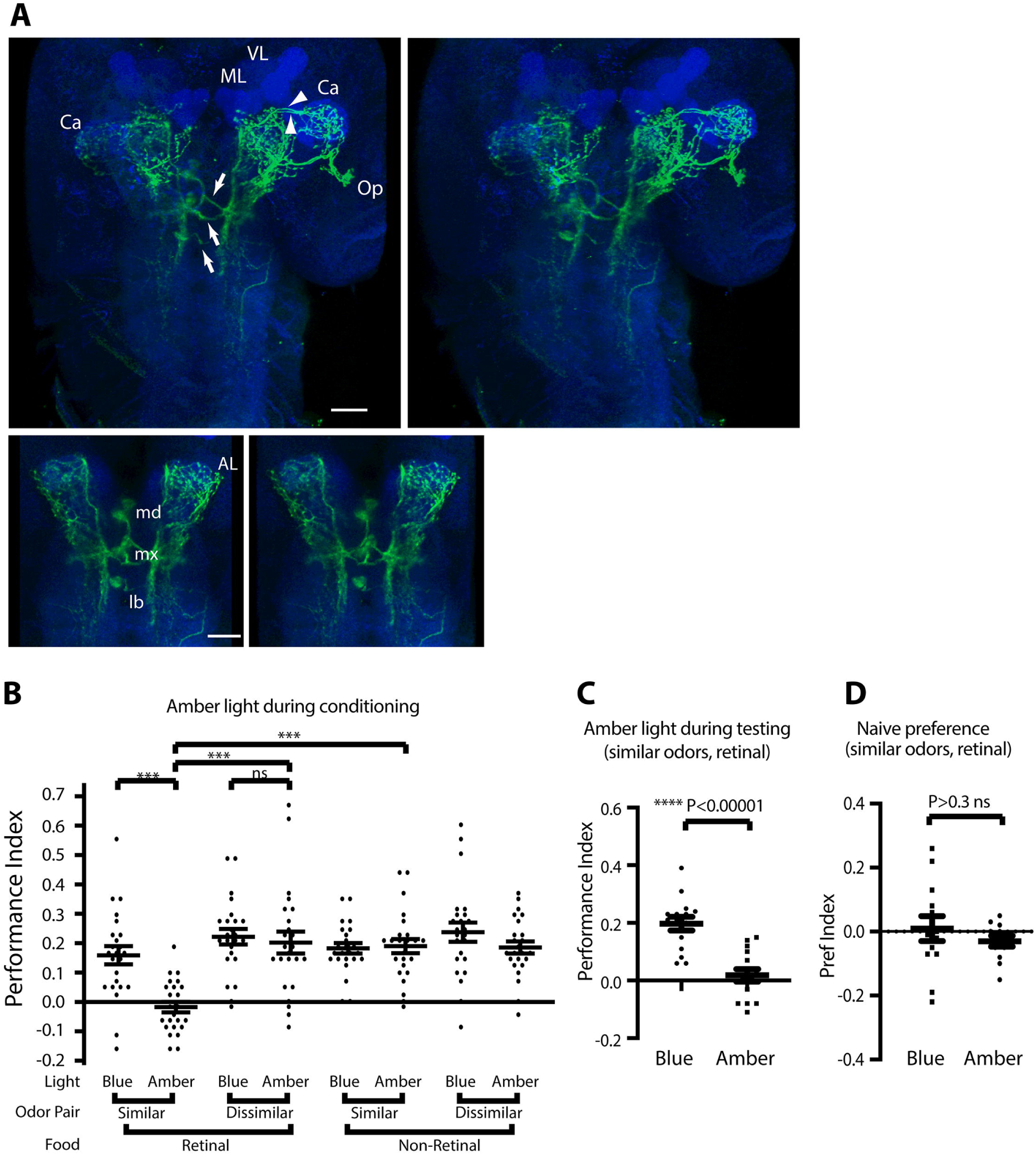
Activation of a small subset of OA neurons including the sVUM1 neurons disrupts odor discrimination but not learning. **A**. Third-instar larvae of genotype *w, FRT*.*GAL80*; *Tdc2-LexA(II) / LexAop2-FLPL(II)*; *GMR34A11-GAL4(III)* / *UAS-CsChrimson*.*mVenus(III)*, used for behavior, generated as in Fig. 4A. Top panels show a pair of stereo images with CsChrimson.mVenus expressed in a subset of 5 sVUM neurons, including calyx-innervating sVUM1 neurons. Arrows indicate the secondary processes of the 3 sVUM1 clusters at the midline. The two tracts entering the calyx are indicated by arrowheads. Scale bar, 40 µm. The bottom row shows a close-up pair of stereo images of the vicinity of the SEZ; 2 cell bodies of sVUM1md cluster, 2 cell bodies of sVUM1mx cluster, and one cell body of the labial cluster are labeled. Antennal lobe (AL), calyx (Ca), optic lobe (Op), MB medial (ML) and vertical (VL) lobes act as landmarks. Scale bar, 20µm. **B**. Activation of the neurons labeled in **A** by CsChrimson in the presence of retinal and amber light (applied during conditioning), abolishes odor choice learning using a similar odor pair, compared to controls exposed to blue light, lacking retinal, or tested with a dissimilar odor pair. No effect of CsChrimson activation is seen on learning using a dissimilar odor pair. Planned statistical comparisons shown are t-tests (***, P < 0.0001, NS, P > 0.6, n=24). An ANOVA test of all data from two experimenters showed no significant effect of experimenter alone, or as interactions with other factors (P>0.2). ANOVA tests showed no significant effect of odor alone (P>0.03, not significant with multiple testing), retinal alone (P>0.4), or light alone (P>0.2), when comparisons that would have included the retinal, amber, similar combination with the strong effect were excluded. **C**. Amber light abolishes learning using a similar odor pair in the presence of retinal, if applied only during testing, after conditioning in blue light (P<10^−5^, t-test, n=15). A two-way ANOVA showed no significant difference between two experimenters (P>0.25), nor any interaction between experimenter and light (P>0.7). **D**. No effect of light color on naive odor preference. Larvae of the same genotype, grown on retinal-containing food as in **B** and **C**, were tested for naive odor preferences under either blue or amber light. No effect was detected (paired t-test, n=12).

Activation of these OA neurons by amber light, during conditioning, had no effect on the ability of larvae to discriminate odors in an appetitive odor discrimination learning assay using a dissimilar odor pair; however, it abolished their ability to discriminate a similar odor pair, suggesting that odor discrimination is affected by activation of these neurons, but not underlying learning ability (Fig. 9B). To exclude any effects of OA activation on the unconditioned stimulus pathway during differential conditioning using the similar odor pair under amber light, we tested for an effect of amber light applied during testing, after conditioning as normal in blue light. Similar to activation during conditioning (Fig. 9B), CsChrimson activation during testing also abolished odor discrimination learning that was seen under blue light (Fig. 9C). Underlying odor preferences between the similar odor pair were also not affected by OA neuron activation, nor by the amber light used to activate them (Fig. 9D).

## Discussion

### OA neurons target extrinsic neurons within the calyx circuitry

Two OA neurons originating in the SEZ, sVUMmd1 and sVUM mx1, innervate the same brain neuropiles, with postsynaptic processes in the SEZ and presynaptic processes in the antennal lobe and MB calyces (Fig. 2). Using GRASP, we found contacts of sVUM1 presynaptic terminals with KCs, PNs, Odd and APL neurons in the calyx. Most of these overlap with OA boutons, suggesting that sVUM1 terminals are mainly presynaptic, acting on presynaptic regions of PNs and the APL, and dendritic regions of KCs and the Odd neurons. Most third-instar KCs have about 6 dendritic processes ending in a claw around a calyx glomerulus (Masuda-Nakagawa et al. 2005), and our GRASP counts of sVUM1-KC contacts overlapping with OA are far fewer than the termini that would be needed to synapse onto the 250-300-KCs present in third-instar larvae (Pauls et al. 2010). Eichler et al. (2017) also find inputs of either sVUM1 neuron (which they call OAN-a1 and OAN-a2) into fewer than 10-15% of KCs in first-instar larvae. Therefore, context-dependent signaling by OA in the calyx must principally affect MB activity via other MB neurons, rather than by direct action on KC dendrites, although the sVUM1 neurons may act directly on a subset of KCs.

Connectomic analysis of a six-hour first-instar larva shows that the sVUM1 neurons, OAN-a1 and OAN-a2 neurons (Eichler et al. 2017), with only 28 presynaptic termini marked in a left brain and 39 in a right brain (Fig. 5), have a qualitatively similar but less extensive calyx innervation pattern than we observe in third instar, with around 89 OA-positive boutons per calyx, and even more active zones, assuming multiple active zones per bouton. Neuromodulatory inputs might develop later in development, as they might need experience-dependent activity to develop, and when behavioral demands increase at a more mature state. Processes of sVUM1 neurons throughout the calyx were more elaborated in the third instar (Fig. 1) compared to the sparse branching of sVUM1 neurons in the first-instar larval connectome image (Fig. 5). Consistent with our findings, Eichler et al. (2017) also report presynaptic contacts of sVUM1 neurons with Odd neuron dendrites, and some synapses of sVUM1 neurons with KCs, but not with most KCs; they do not comment on synapses with PNs or the APL.

### Roles of APL and Odd neurons in calyx activity

sVUM1 presynaptic termini make many contacts with both the APL and Odd neurons in third-instar larvae (Fig. 3, 4). Odd neurons ramify throughout the calyx and receive input from PNs generally, potentially forming a channel for non-selective odor processing that is parallel to the main MB odor-specific processing through KCs. sVUM1 neurons could potentially change the Odd neuron gain or tuning properties, to signal changes in behavioural state that guide odor-driven choice behaviours, for example during chemotactic behaviour, in which Odd neurons are implicated (Slater et al. 2015).

The APL mediates a negative feedback loop from KC outputs in the MB lobes to KC inputs in the calyx, thus potentially both limiting the duration of KC activity and improving their odor discrimination (Masuda-Nakagawa et al. 2014; Lin et al. 2014). sVUM1 synapses onto the APL in the calyx could therefore potentially regulate this feedback loop. This could increase signal-to-noise ratio, in a context-dependent manner, by sharpening odor representations in the calyx via APL inhibitory feedback, similar to the “gain control” mechanism with enhancement of behaviorally relevant responses and suppression of non-relevant ones in monkey visual system (Treue and Martínez Trujillo 1999; Gilbert and Li 2013). On the other hand, at extremes, this would also decrease the sensitivity to input, and hence decrease learning. Inhibition of the APL enhances learning, by increasing the sensitivity to input (Liu and Davis 2009).

In addition, since we observed some GRASP signals adjacent to GABA termini (Fig. 3E), APL feedback could also inhibit OA release from sVUM1 termini, further increasing the complexity of interactions between OA innervation and the KC/APL negative feedback loop. NA regulation of inhibitory neurons is also a feature of the mammalian olfactory circuitry. In the olfactory bulb, disinhibition of mitral cell (equivalent to PN) activity by NA regulation of inhibitory granule cells has been proposed (Nai et al. 2009). In mammalian PCx, feedforward and feedback inhibition are postulated to enhance cortical representation of strong inputs (Stokes and Isaacson 2010), and the PCx receives extensive NA innervation from the LC, although its role in modulating inhibition has not been investigated.

### PNs as potential targets of OA modulation

GRASP analyses suggest that PNs are postsynaptic to the sVUM1 neurons (Fig. 3). Indeed, an Oamb::eGFP exon-trap fusion is localized on PN presynaptic terminals, albeit more widely than GRASP puncta (Fig. 6). Similarly, the honeybee Oamb ortholog AmOA1 is found widely in the calyx (Sinakevitch et al. 2011), although these authors do not distinguish between PN terminals or KC dendrites. Much aminergic neurotransmission acts via extrasynaptic receptors (Bentley et al. 2016), and this may be the case for Oamb in PN terminals. sVUM1 neurons have dense core vesicles (DCV), and these might release peptides together with OA (Tao et al. 2019). However, it is known that OA is packed into vesicles by DVMAT-A (Greer et al. 2005) in *Drosophila*, and although we can not exclude the possibility of volume transmission rather than classical synapses as an extra mechanism of release, this has not been documented for OA.

Oamb is a GPCR that signals apparently through Gq, to release Ca^2+^ from intracellular stores (Balfanz et al. 2005; Morita et al. 2006); it may also elevate cAMP (Han et al. 1998), although this effect appears smaller (Balfanz et al. 2005). In adult *Drosophila*, Tomchik and Davis (2009) found that bath application of OA increased cAMP responses in PN axons. These results suggest that OA can excite PN neuron terminals via cAMP, and thus facilitate synaptic transmission. In agreement with this, presynaptic facilitation was observed in *Aplysia* sensory neurons through the cAMP/PKa pathway after exogenous expression of the OA receptor, Ap oa_1_ (Chang et al. 2000).

Since we also observe GRASP signals between sVUM1 neurons and a minority of KCs, a cAMP mediated mechanism in KCs could also facilitate synaptic transmission; elevating cAMP facilitates Ca^2+^ responses of MB neurons to stimulation by ACh (Tomchik and Davis 2008). The cAMP/PKA pathway in MB neurons can also affect the duration of excitation through a K^+^ channel mediated mechanism (Aoki et al. 2008). However, we did not observe Oamb receptor expression in KC dendrites.

An alternate mechanism of OA in plasticity is that subthreshold sensory input could be gated by OA to facilitate the detection of subthreshold signals. In the mammalian olfactory bulb, LC input facilitates the detection of peri-threshold stimuli and near-threshold rewarded odors (Jiang et al. 1996; Escanilla et al. 2012), via an increase in mitral cell excitability mediated by NA action on α1-adrenergic receptors (Ciombor et al. 1999; Hayar et al. 2001). Therefore, Oamb receptors in PN terminals in the calyx could potentially participate in plastic changes to facilitate the detection of behaviorally relevant sensory input in a given behavioral context.

### Odor discrimination learning

We selected odorants as a dissimilar pair, EA and PA, that activate different sets of glomeruli in the AL (Kreher et al. 2008). On the other hand, odor mixtures, which we used as the basis of our similar odor pairs, often activate patterns in the olfactory centres that are a combination of single odorant response in honeybees (Joerges et al. 1997) and mice (Grossman et al. 2008). How this pattern of activity is translated into perceptual similarity is not entirely clear, but representations in the cortex do correlate with behavioural discrimination (Chapuis and Wilson 2012). Reinforced olfactory discrimination has been used in honeybees (Stopfer et al. 1997), *Drosophila* (Lin et al. 2014) and mammals (Linster and Cleland 2001), to test neural mechanisms of odor discrimination.

Here we observed that optogenetic activation of 5 neurons, including the VUM1 neurons, compromised discrimination of similar odors in an appetitive conditioning paradigm, either during conditioning, or during testing (Fig. 9). This result is consistent with OA innervation affecting the selectivity of odor representations by KCs, both during formation of odor memory, and during recall. One explanation is that VUM1 activation enhanced the gain of stimulus-driven PNs, increasing the magnitude and number of KCs responding, and therefore making odor representations by KCs overlap more between similar odors, and thus lowering discriminability of similar odors. Another possibility is that the detection of odors is compromised as a result of inhibition of stimulus induced activity, perhaps by action of VUM1s on the GABAergic APL.

A role for OA as a reinforcer in appetitive associative learning has been shown in honeybees and flies. The honeybee VUMmx1 neuron has properties of a reinforcer; its depolarization could replace sugar reinforcement in appetitive learning (Hammer 1993), and injection of OA into the MB calyx induced memory consolidation (Hammer and Menzel 1998). On the other hand, it has been proposed that VUMmx1 learns about the value of the odor, and acts as a prediction error signal in appetitive learning (Menzel 2012). However, associative plasticity in the MBs in *Drosophila* is thought to reside mainly in the lobes rather than the calyx, for both appetitive (Schwaerzel et al. 2003; Schroll et al. 2006; Liu et al. 2012) and aversive learning (Aso et al. 2014, 2012). OA as a reinforcer in appetitive learning appears to act via Oamb expressed in PAM dopamine neurons that target the medial MB lobes. The *NP7088-GAL4* line that includes the fly equivalent of the honeybee VUMmx1, OA-VUMa2, did not induce and is not required for appetitive learning in adult *Drosophila* (Burke et al. 2012). In larvae, pPAM neurons that innervate the medial lobe appear to be involved in reward learning (Rohwedder et al. 2016). We found Oamb::GFP at the tip and around the MB medial lobe, suggesting this as a potential site of integration of appetitive reinforcement for learning. Furthermore, Oamb is required in KCs for adult appetitive learning (Kim et al. 2013), suggesting some direct input of an OA-encoded appetitive signal into KCs; the localization of Oamb to MB lobes (Crittenden et al. 1998) is consistent with OA signaling on KC axons in the lobes rather than dendrites in the calyx. Taken together, OA action as a reinforcer in associative learning might occur via unidentified inputs into dopaminergic neurons or KC lobes.

sVUM1 neurons innervate the AL, and synapse with inhibitory interneurons (Supplementary File S1), therefore OA can regulate the processing of olfactory signals. In honey bee injection of OA in the antennal lobes impairs memory acquisition and recall but not odor discrimination (Farooqui et al. 2003). Noradrenalin in mammals has a role in olfactory memory formation by affecting the inhibitory network in the olfactory bulb (reviewed by Kaba and Nakanishi 1995). Therefore, OA is likely to be involved in synaptic plasticity in the first relay of the olfactory pathway, sharpening its output. In our studies we tested odor discrimination ability, and our results favor the interpretation that OA acts by changing the efficacy of synaptic input to the MBs, at the calyx. A role in gating behaviorally relevant sensory input, is favored by our behavioral data, and is also suggested as a role of OA in regulating the threshold response of peripheral sensory receptors and afferents in insects and NA in the CNS of mammals (Berridge and Waterhouse 2003). Our behavioral data could be interpreted to result from an increase in the levels of sensitivity, at expense of discrimination, to meet the demands of changes in behavior, signaled by OA.

### Perspectives

Sensory representations are dynamically modified by higher brain signaling, according to behavioral states such as attention, expectation, and behavioral task; and LC (locus coeruleus) activation in mammals and OA activation in insects correlate with changes in behavioral states. Mammalian olfactory neuropiles are densely innervated by noradrenergic input, similar to the dense innervation of the AL and calyx by OA sVUM1 neurons in insects. The innervation of sVUM1 neurons throughout the calyx, and their potential synaptic connections to PNs, KCs, APL, and Odd-like neurons, mean that OA could induce a network level switch either by gating input afferent activity, and/or by interacting with the KC/APL feedback loop, and thus also affecting the output activity from the calyx – not only through KCs but also potentially via the Odd neurons. Behavioral demands would determine the balance between sensitivity and discrimination via OA; whether to escape from a predator at all cost, or the need for fine discrimination to recognize food.

## Materials and Methods

### Genetics and molecular biology

#### Fly Stocks

Flies were raised on standard cornmeal medium at 25°C and subjected to a 12 hour day/night cycle. Stocks used are listed in Table 1.

**Table 1.**
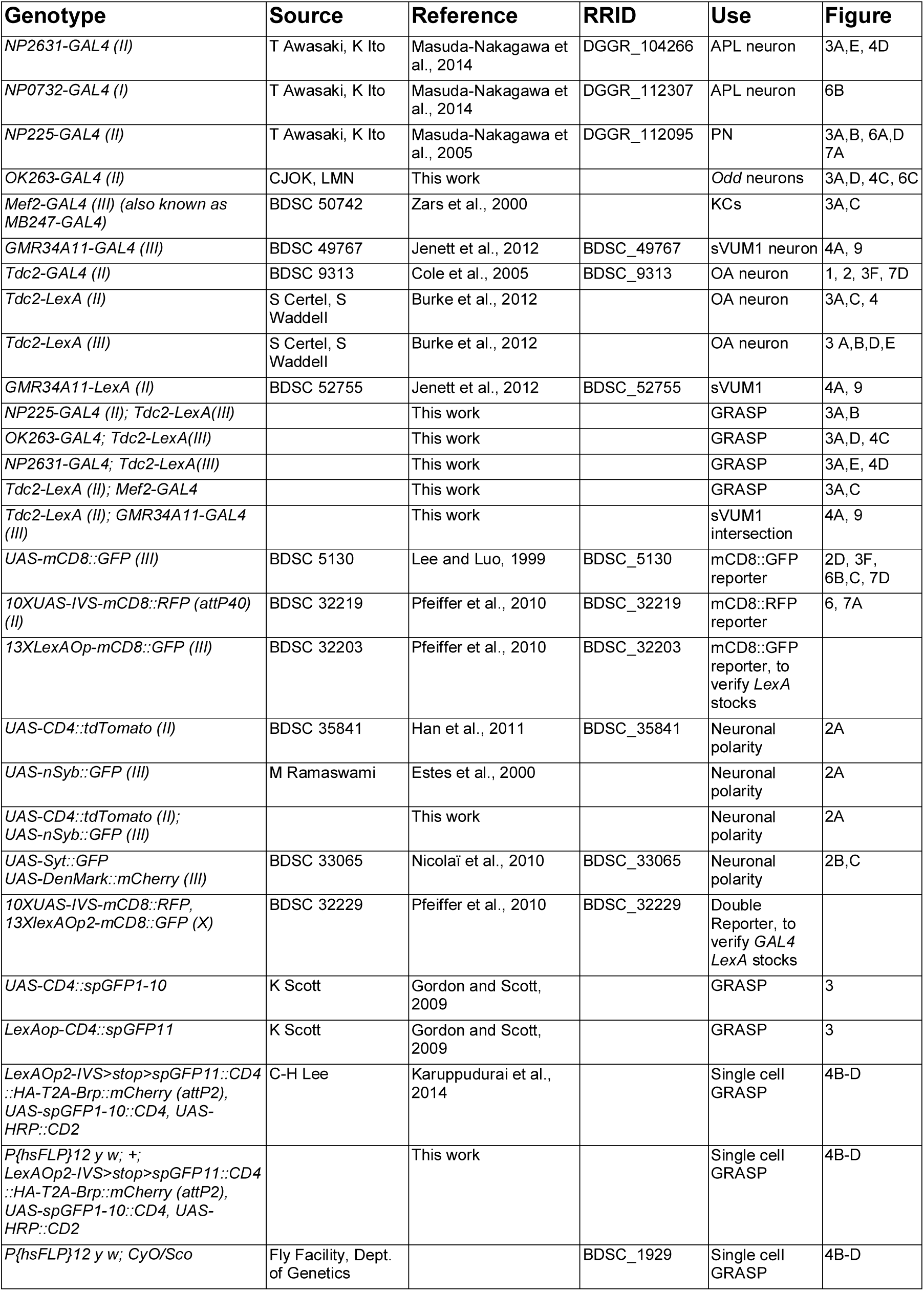
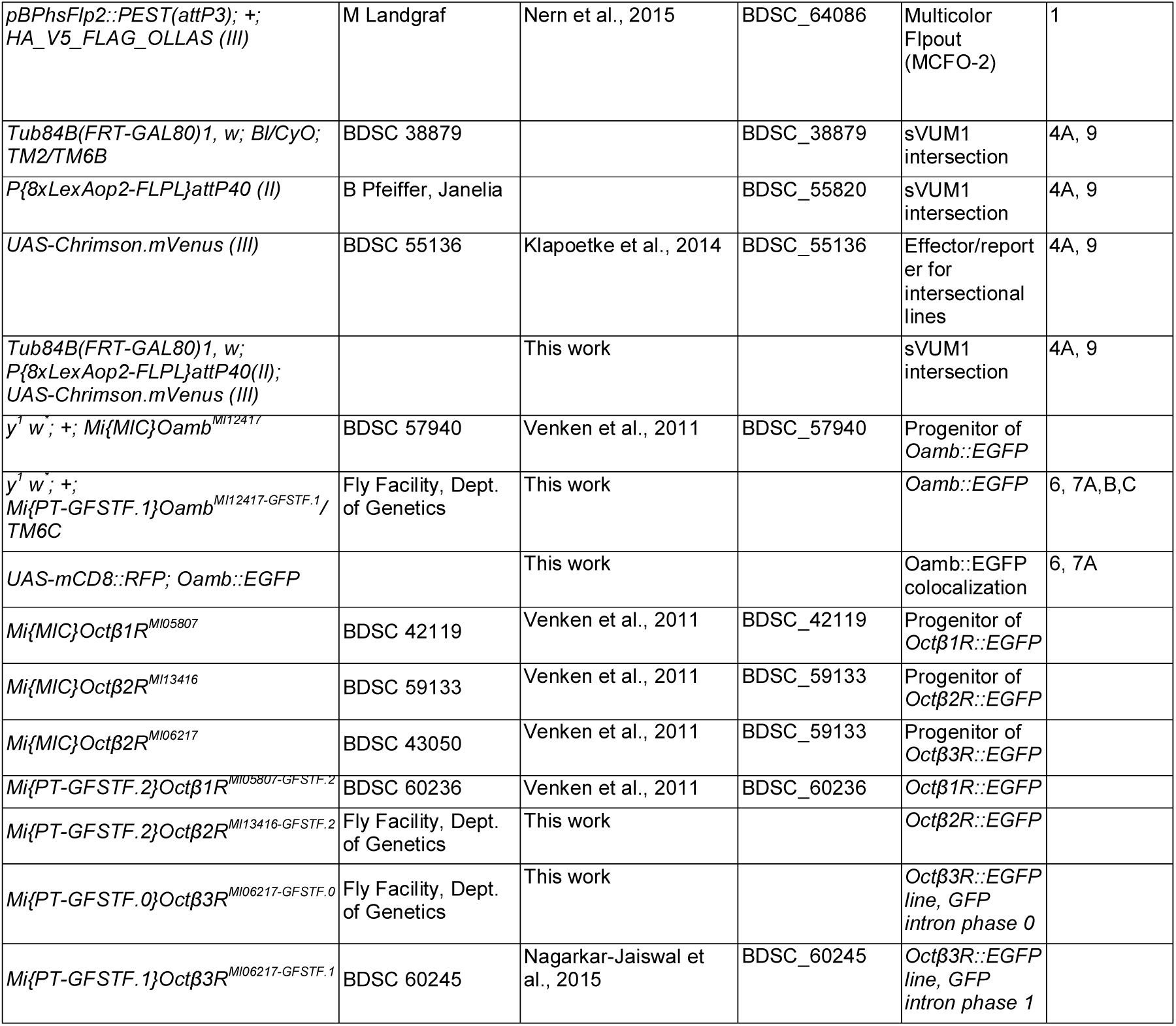
*Drosophila* stocks used. Includes all genotypes used for this study, including those that do not appear in figures.

#### MultiColor FlpOut

MultiColor FlpOut was performed according to Nern et al. (2015). Females of genotype *pBPhsFlp2::PEST(attP3); +; HA_V5_FLAG_OLLAS* (“MCFO-2”) were crossed with male *Tdc2-Gal4* flies. Parents were left in vials to lay eggs for 24-hour intervals. After another 24 hours, larval progeny were heat shocked at 35-37°C by immersion in a heated circulated waterbath for 15-30 minutes, thus ensuring larvae were aged 24-48 hours after egg-laying (AEL) at the time of heat shock.

#### GRASP

Standard GRASP was according to Gordon and Scott (2009). A line carrying GRASP constructs *UAS-CD4::spGFP1-10* and *LexAop-CD4::spGFP11*, was crossed to individual *LexA GAL4* lines as needed: *NP225-GAL4(II); Tdc2-LexA(III)*, or *OK263-GAL4(II); Tdc2-LexA(III)*, or *NP2631-GAL4(II); Tdc2-LexA(III)*, or *Tdc2-LexA(II); Mef2-GAL4(III)* (Fig. 3) or to individual *GAL4* or *LexA* lines as controls (Fig. S1). Reconstituted GFP was detected using rat monoclonal anti-GFP. This did not detect either of the GRASP components GFP1-10 or GFP11, when *Gal4* or *LexA* drivers were used alone (FigS1). GRASP signals had to meet a criterion of occurring in two consecutive 0.5-µm confocal sections. For single cell GRASP (Karuppudurai et al. 2014), we generated larvae carrying *P{hsFLP}*12, appropriate *GAL4* and *LexA* combinations, and a recombinant chromosome with insertions *LexAOp2-IVS>stop>spGFP11::CD4::HA-T2A-Brp::mCherry (attP2), UAS-spGFP1-10::CD4*, and *UAS-HRP::CD2*, by generating a stock carrying *P{hsFLP}12 y w; +; LexAOp2-IVS>stop>spGFP11::CD4::HA-T2A-Brp::mCherry (attP2), UAS-spGFP1-10::CD4, UAS-HRP::CD2*, and crossing females of this stock to relevant *GAL4 LexA* males. To generate labelled single cells, parents were allowed to lay eggs initially for 24-hour intervals, then for 6-hour intervals in vials containing half the amount of food. At 0-24 h, 24-48 h, or later at 12-18 h, 18-24 h, or 24-30 h AEL, progeny were heat shocked as above for 10-50 minutes at 37 □C. Progeny were incubated from RT until dissection of non-tubby wandering third-instar larvae.

#### Generation of an EGFP-tagged Oamb line

The *Mi{MIC}Oamb*^*MI12417*^ insertion in coding intron 3 of *Oamb* at 3R:20697059, (BDSC stock 57940; henceforth referred to as *MI12417*) was verified by PCR using primers *MI12417-5F/MiMIC-5R1* for the 5’ end and *MI12417-3R/MiMIC-3F1* for the 3’ end (Table 2; Supplementary Fig. 2). Sequencing of these PCR products and alignment with the *Drosophila* genome sequence using BLASTN (Altschul et al., 1990; https://blast.ncbi.nlm.nih.gov/) showed insertion of *MiMIC* at the recorded site of 3R 20697058-9 (Supplementary Figs. 3,4). The location of the *MI12417* insertion site relative to *Oamb* coding exons was determined by using Oamb-B sequences for BLASTN and TBLASTN queries of the *Drosophila* genome assembly (http://flybase.org/blast/; Supplementary Fig. 5). TMHMM (Sonnhammer et al., 1998; http://www.cbs.dtu.dk/services/TMHMM/) was used to predict the amino-acid coordinates of Oamb transmembrane (TM) domains (Supplementary Figs. 6,7).

**Table 2.**
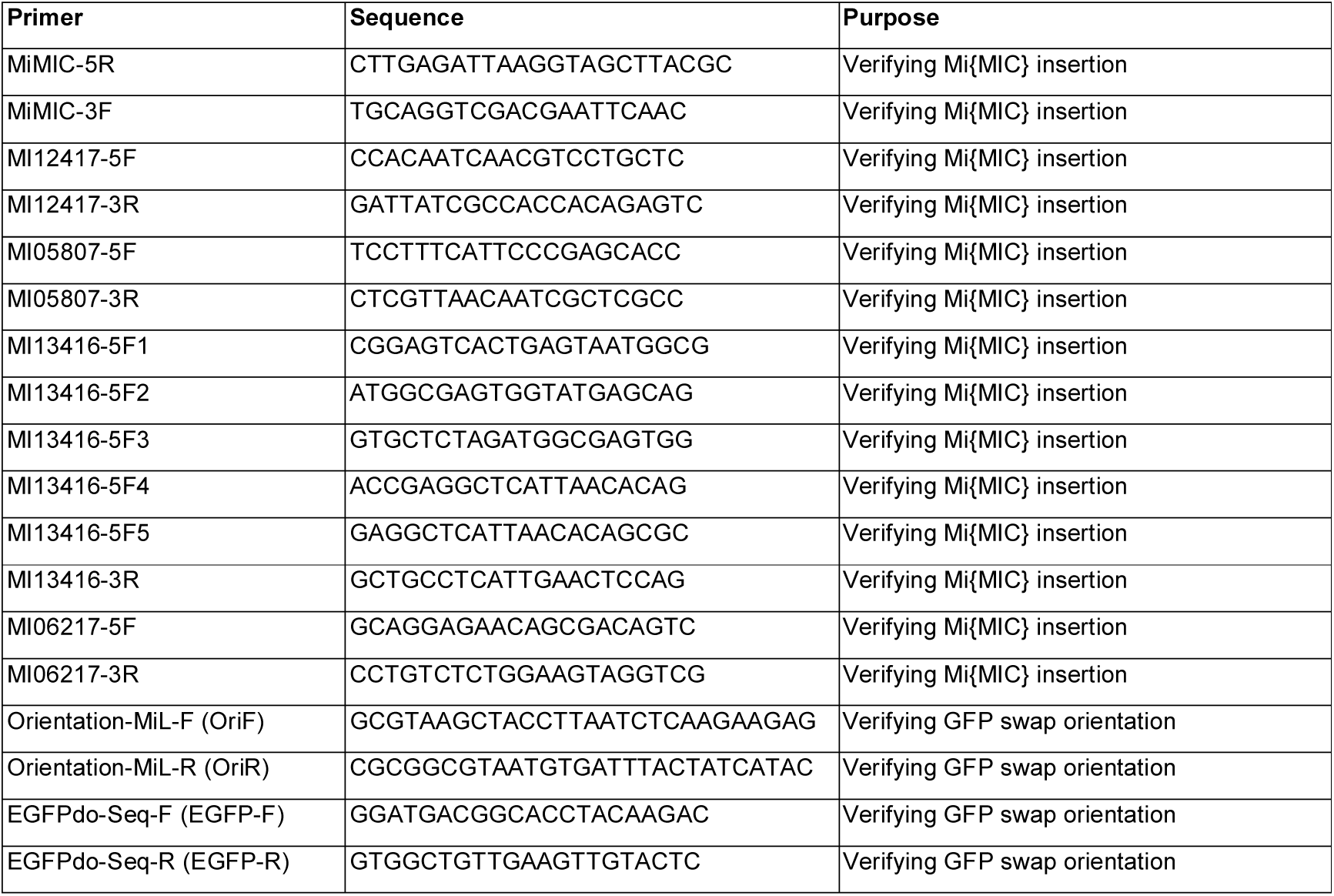
Primers.

To insert an EGFP-encoding exon into the *MI12417* insertion by RMCE, we chose the splice phase-1 version of the *EGFP-FlAsH-StrepII-TEV-3xFlag* plasmid (DGRC 1306; Venken et al. 2011) as recommended by the Baylor Gene Disruption Project (http://flypush.imgen.bcm.tmc.edu/pscreen/rmce/rmce.php?entry=RM00888). This was co-injected with a helper phiC31-integrase plasmid (Venken et al. 2011) by the *Drosophila* microinjection facility (Department of Genetics, University of Cambridge). Injected embryos were left to hatch into adult flies and crossed to a *y w* double balancer stock. RMCE events were identified by loss of the *MiMIC yellow*^*+*^ marker in F1 progeny. Four PCR reactions were carried out to determine the orientation of the EGFP cassette in each recombinant *Oamb::EGFP* stock (Table 2) as described in Venken et al. (2011).

#### *OctβR* EGFP fusions

For *Octβ1R* and *Octβ3R*, we used lines *Octβ1R*^*MI05807-GFSTF*.*2*^ and *Octβ3R*^*MI06217-GFSTF*.*1*^ (Bloomington stocks 60236 and 60245), respectively, as *Octβ1R::EGFP* and *Octβ3::EGFP* exon traps. For *Octβ2R*, we used *Mi{MIC}Octβ2R*^*MI13416*^ to generate *Octβ2R*^*MI13416-GFSTF*.*2*^ using a similar approach as that described above for Oamb::EGFP. We also generated a second *Octβ3::EGFP* exon trap, *Octβ3R*^*MI06217-GFSTF*.*0*^, in a different reading frame from Bloomington stock 60245, to allow for the possibility of the *EGFP* exon being spliced as in transcripts *RJ* or *RK* (frame 0, with only *RJ* able to encode all seven TM domains), rather than *RF* or *RG* (frame 1). Positions of each insertion were confirmed by PCR and sequencing similarly to Oamb, using primers as described in Table 2.

#### Molecular methods

Genomic DNA was extracted from 15-30 flies (1-7 days after eclosion) and homogenized in 100 mM Tris-HCl, 8.5; 80 mM NaCl, (Sigma 31434); 5% Sucrose (Sigma S0389); 0.5% SDS (Sigma L4509); 50 mM Na-EDTA (Sigma ED2SS), pH 8.0. The homogenate was incubated with RNase A (Roche 10109142001) for 1 hour at 37°C followed by Proteinase K (Roche 03115887001) for 1 hour at 50°C, and purified with phenol-chloroform (Sigma 77617) and chloroform (Sigma C2432). DNA was precipitated with 0.6 volumes of isopropanol (Sigma, 59304) and washed with 75% ethanol (Sigma E7023), dried overnight at room temperature and re-suspended in 10 mM Tris-HCl pH 8.0 (Sigma T6066).

PCR reactions (20 μl) contained 0.4 μl or 1 μl genomic DNA, 1 μl of each 10 μM primer (Sigma), 2μl of 10X PCR buffer (Qiagen 203203), 0.4 μl of 10 μM dNTP mix (Roche 11581295001), 0.08 μl of 5 U/μl HotStarTaq DNA polymerase (Qiagen 203203) and 15.1 μl or 14.5 μl milliQ water. PCR cycling in a G-Storm Thermal Cycler (GS4) was: 15 minutes at 95°C; 40 cycles of: denaturation at 94°C for 30s, annealing at 60°C for 30s and elongation at 72°C for 1 min; and a final elongation step at 72°C for 10 minutes. PCR products were loaded with 6X DNA gel-loading dye (ThermoFisher R0611) on a 1% Agarose Gel (LifeTech 16500500; 1X TBE buffer, LifeTech 16500500) with GelRed (Biotium 41003-T) for gel electrophoresis. 100 bp DNA ladder was used as a marker (LifeTech 15628019). PCR products were purified using the Qiaquick PCR Purification Kit (Qiagen, 28104), and sequenced at the Department of Biochemistry Sequencing Facility (University of Cambridge).

### Immunohistochemistry and Confocal imaging

Third-instar wandering larval brains (144-176 hours AEL) were dissected in cold PBS (Sigma P4417), fixed in 4% Formaldehyde (Polysciences 18814) / PEM buffer (0.1 M PIPES, Sigma P1851; 2 mM EGTA, Sigma E3889; 1 mM MgSO_4_; NaOH) for 2 hours at 4°C, washed for 3×10 minutes (or 4×15 minutes) in 0.3% Triton-X (Sigma T8787) in PBS (PBT) and incubated in 10% NGS (Normal goat serum; Vector Labs S-1000) in 0.3% PBT for 1 hour at room temperature. Brains were incubated in primary antibody in 10% NGS-0.3% PBT at 4°C for 2-3 days on a mini disk rotor (Biocraft, BC-710), washed for 3×15 minutes with 0.3% PBT and further incubated in secondary antibody in 10% NGS at 4°C for 2-3 days again on the mini disk rotor. Brains were finally washed 1×15 minutes with PBT, followed by 3×10 minutes with PBS, and left in 50% Glycerol/PBS at 4°C for at least one night prior to imaging.

Primary and secondary antibodies are listed in Table 3. Brains were incubated in primary antibody at 4°C for 2-3 nights, washed three times in PBT for 15-minutes, and incubated in secondary antibody for 2-3 more nights. To reduce background with the polyclonal chicken anti-GFP (Abcam, Ab13970), it was pre-incubated with *MI12417* larval brains which do not express GFP. Fifty *MI12417* larval brains were incubated in 1:20 chicken anti-GFP in 10% NGS in 0.3% PBT at 4°C overnight. A further 50 *MI12417* larval brains were added and further incubated at 4°C over 2 nights. Mounting and orientation of brains for image acquisition was as described in the supplemental information in Masuda-Nakagawa et al. (2009). Imaging was carried out using a Zeiss LSM710 Confocal Microscope with a 40X NA1.3 oil objective. Images were processed using ImageJ software (https://imagej.nih.gov/ij/download.html).

**Table 3.**
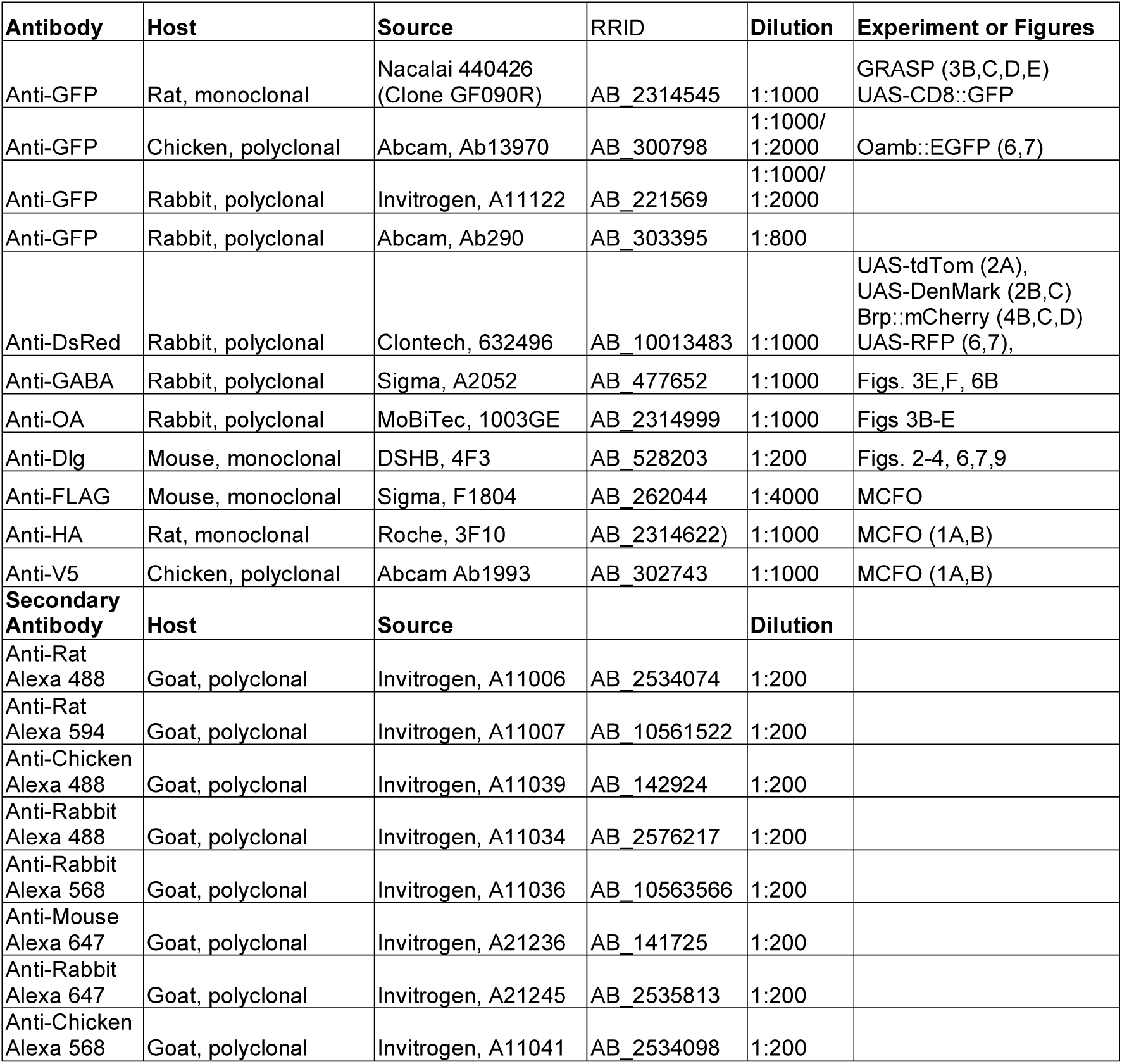
Antibodies.

| Antibody | Host | Source | RRID | Dilution | Experiment or Figures |
| --- | --- | --- | --- | --- | --- |
| Anti-GFP | Rat, monoclonal | Nacalai 440426 (Clone GF090R) | AB_2314545 | 1:1000 | GRASP (3B,C,D,E)<br>UAS-CD8::GFP |
| Anti-GFP | Chicken, polyclonal | Abcam, Ab13970 | AB_300798 | 1:1000/<br>1:2000 | Oamb::EGFP (6,7) |
| Anti-GFP | Rabbit, polyclonal | Invitrogen, A11122 | AB_221569 | 1:1000/<br>1:2000 |  |
| Anti-GFP | Rabbit, polyclonal | Abcam, Ab290 | AB_303395 | 1:800 |  |
| Anti-DsRed | Rabbit, polyclonal | Clontech, 632496 | AB_10013483 | 1:1000 | UAS-tdTom (2A),<br>UAS-DenMark (2B,C)<br>Brp::mCherry (4B,C,D)<br>UAS-RFP (6,7), |
| Anti-GABA | Rabbit, polyclonal | Sigma, A2052 | AB_477652 | 1:1000 | Figs. 3E,F, 6B |
| Anti-OA | Rabbit, polyclonal | MoBiTec, 1003GE | AB_2314999 | 1:1000 | Figs 3B-E |
| Anti-Dlg | Mouse, monoclonal | DSHB, 4F3 | AB_528203 | 1:200 | Figs. 2-4, 6,7,9 |
| Anti-FLAG | Mouse, monoclonal | Sigma, F1804 | AB_262044 | 1:4000 | MCFO |
| Anti-HA | Rat, monoclonal | Roche, 3F10 | AB_2314622) | 1:1000 | MCFO (1A,B) |
| Anti-V5 | Chicken, polyclonal | Abcam Ab1993 | AB_302743 | 1:1000 | MCFO (1A,B) |
| **Secondary Antibody** | **Host** | **Source** |  | **Dilution** |  |
| Anti-Rat Alexa 488 | Goat, polyclonal | Invitrogen, A11006 | AB_2534074 | 1:200 |  |
| Anti-Rat Alexa 594 | Goat, polyclonal | Invitrogen, A11007 | AB_10561522 | 1:200 |  |
| Anti-Chicken Alexa 488 | Goat, polyclonal | Invitrogen, A11039 | AB_142924 | 1:200 |  |
| Anti-Rabbit Alexa 488 | Goat, polyclonal | Invitrogen, A11034 | AB_2576217 | 1:200 |  |
| Anti-Rabbit Alexa 568 | Goat, polyclonal | Invitrogen, A11036 | AB_10563566 | 1:200 |  |
| Anti-Mouse Alexa 647 | Goat, polyclonal | Invitrogen, A21236 | AB_141725 | 1:200 |  |
| Anti-Rabbit Alexa 647 | Goat, polyclonal | Invitrogen, A21245 | AB_2535813 | 1:200 |  |
| Anti-Chicken Alexa 568 | Goat, polyclonal | Invitrogen, A11041 | AB_2534098 | 1:200 |  |

**Table 4.** Other reagents.

| Reagent | Source | Experiment or Figures |
| --- | --- | --- |
| Paraffin oil | Sigma-Aldrich, 76235 | Behavior |
| Pentyl Acetate | Sigma-Aldrich, 109584 | Behavior |
| Ethyl Acetate | Sigma-Aldrich, 319902 | Behavior |
| Ethanol | Sigma-Aldrich, 32221 | Behavior |
| all-trans-retinal (ATR) | Sigma-Aldrich, R2500 | Behavior |
| D-(-)-Fructose | Sigma-Aldrich, 47740 | Behavior |
| Agarose | Sigma-Aldrich, A9539 | Behavior |

### Behavioral assay

#### Larval culture

Males of genotype *w; Tdc2-LexA; GMR34A11-GAL4/TM6B* were crossed to females of genotype *Tub84B(FRT-GAL80)1, w; LexAop-FLP; UAS-Chrimson*.*mVenus/TM6B* to generate F1 larvae in which *UAS-Chrimson*.*mVenus* could be expressed only in cells expressing both *Tdc2-LexA* and *GMR34A11-GAL4*, in which LexA-dependent *FLP* expression had removed the GAL4 inhibitor GAL80. Larvae were allowed to develop in food vials containing 200 µM all-trans-retinal (Sigma, R2500), in the dark at 21 degrees. For both the retinal and non-retinal food vials, transfer of adults into new vials was performed both in the morning and in the evening, to then collect them after 108-120 hours in the case of non-retinal vials at 25°C, and after 132-144 hours for those kept in retinal vials at 23°C.

#### Behavioral arena

8.5-cm petri dishes containing agarose were prepared the day before use, using 100 ml of distilled water with 0.9% agarose (Sigma A9539). Fructose petri dishes were prepared similarly, but containing 35% fructose (Sigma-47740). Petri dishes had perforated lids to facilitate creation of odorant gradients within the dish, generated by sucking air away using a benchtop fume extractor (Sentry Air Systems, SS-200-WSL) at the back of the assay platform. Odorants were diluted in paraffin oil (Sigma-Aldrich 76235), and 10-µl aliquots were pipetted with a cotton filled tip, immediately before conditioning or test, into custom-built Teflon containers with pierced lids (7 holes), made in the Physiology workshop in the University of Cambridge, based on samples kindly provided by B. Gerber.

#### Light apparatus

Our optogenetics apparatus was constructed as described by de Vries and Clandinin (2013) for activation of ChR. We modified it to shine amber light to activate CsChrimson. A BK Precision 1698 Model DC power pack was connected to a pulse generator, driving 4 sets of amber light LEDs (591 nm), Luxeon star Amber LED on Tri-Star Base, 330 lm at 350mA (Cat No. SP-03-A5). The irradiance on the platform was 0.06 µW/mm^2^ on average on the 8.5 cm plate (24 µW on a 20×20mm sensor). The pulse generator was custom built by the Psychology Department workshop of the University of Cambridge, to deliver 10-ms pulses at 10Hz for 30s, followed by 30s without pulses. This cycle was repeated 5 times, making a conditioning step of 5 minutes in total. The power supply was run at 17-mV constant voltage. This pulse frequency and width were chosen to replicate the activity of the only recorded sVUM1 neuron, the honeybee sVUMmx1, when activated by a sucrose reward (Hammer 1993).

#### Behavior conditioning

Third-instar larvae were collected from vials using a stainless steel SANPO µm/m 355 and washed with tap water. Larvae were washed twice by transferring through a drop of tap water, and then placed on the conditioning agarose plate (35% fructose) with the help of a paint brush.

One conditioning cycle consisted of placing larvae in a fructose dish exposed to the odor to be conditioned with for 5 minutes, and then washing in 2 drops of water to clean them of fructose, before transferring to an agarose dish, where larvae were exposed to the other odor of the pair for 5 minutes without the reinforcer. This cycle was repeated 3 times. For experiments involving activation of OA neurons, the 3 conditioning cycles with reinforcer present and the 3 non-reinforced cycles were carried out under pulses of amber light (Fig. 9). For controls without activation of CsChrimson, conditioning was performed as with amber light but under dim blue light, using an aquarium lamp, covered with paper to decrease its intensity.

Containers were placed at 1 cm from the side of the dish (Fig. 8). One container was filled with odor while the one on the opposite side contained mineral oil. One experimenter (ADM) placed containers consistently at the same position throughout conditioning cycles; the other (MM) alternated oil and odor containers between sides for each 5 min of the 3 cycles, to avoid any inadvertent effects of illumination (MM).

#### Odor dilutions

Our choice of odor was based on the cluster analysis of Kreher et al. (2008). We measured the response index (RI) of Canton-S (CS) wild type larvae to diverse odorants of the Kreher odor panel at different intensities, aiming to reach an RI of above 0.5, according to the widely used protocol of Monte et al. (1989), Rodrigues and Siddiqi (1978), and Kreher et al. (2008). RI was defined as RI=(S-C) / (S+C), where S is the number of larvae on the odor side, C is the number of larvae on the diluent side, after 5 minutes.

We selected ethylacetate (EA; Sigma-Aldrich, cat no. 319902) at 1:2000 dilution which gave RI=0.57 ± 0.04 (mean ± SEM, n=12), in CS third-instar larvae. To establish a dissimilar odor pair, we selected pentyl acetate (PA; also known as amyl acetate or n-amyl acetate; Sigma-Aldrich, cat no. 109584). EA and PA are homologous esters that differ by a length of three carbons. Since the RI for EA is higher than PA (Cobb and Dannet 1994), we used odor balancing to determine the dilution of PA that would balance EA, i.e., a preference index (Pref-I) of around zero given a choice between the two odors, first using CS wild type larvae (Supplementary Methods), and subsequently with larvae of the genotype used for learning and discrimination behavior, *w, FRT*.*GAL80*; *Tdc2-LexA / LexAop2-FLPL*; *GMR34A11-GAL4* / *UAS-CsChrimson*.*mVenus*,. Pref-I was calculated as (number of larvae in EA) -(number of larvae in PA) / (total number of larvae) (Scherer et al. 2003). A negative Pref-I means preference towards PA. Dilutions of EA at 1:2000 and PA at 1:500 gave a Pref-I of close to zero, and these were used as dissimilar odors in conditioning experiments.

#### Testing

Larvae were tested by placing them on an agarose plate carrying a container with EA (1:2000) on one side, and a container with PA (1:500) on the opposite side (dissimilar odors). or a container with EA:PA 4:1 and one with EA: PA 1:4 (similar odors; mixes were made using the same dilutions as in the dissimilar odor pair). Testing was performed under dim blue light, as used during conditioning, for 5 minutes. Larvae were counted on the side of the conditioned odor, the unconditioned odor, and in the neutral zone in the middle. A single performance Index (PI) (Selcho et al. 2009) was calculated as:

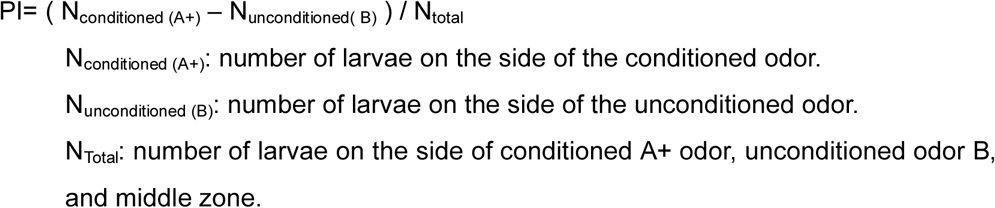

Since we performed a reciprocal conditioning run, to avoid non-associative effects, a different sample of larvae were conditioned in parallel, with the previously unconditioned odor. An average Performance Index (PI) was calculated from the two groups of larvae (A+/B and A/B+) conditioned reciprocally, using the formula:

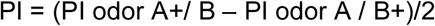

To test the effect of amber light during testing, the same procedure of differential conditioning with similar odors, and larvae grown in retinal-supplemented food, was performed by conditioning in blue light, and then applying either amber or blue light during testing.

#### Controls for the effect of light on similar odor responses

To test for any bias of larval behavior caused by amber light alone, larvae of the experimental genotype were exposed to light, in similar light conditions as in the conditioning trainings, but without fructose, and tested for their odor preferences. Thirty larvae grown in retinal food were exposed to the pair of similar odors on opposite sides of the dish, for 5 minutes, under either blue light or amber light, and their preference index calculated. Containers were swapped in order to exclude bias on illumination. Trials with dissimilar odors were also included,

### Statistical analysis

Single planned comparisons were performed using t-tests in Microsoft Excel. A four-factor ANOVA using SPSS software was used to test whether performance index was affected by food type (retinal vs. non-retinal), light (blue vs. amber), odor pairs (similar pair vs. dissimilar pair), or experimenter (either of two). Effects of experimenter on learning scores were tested using 2-way ANOVA tests in GraphPad Prism 8.0 and found to be non-significant (P>0.2). Planned comparisons between the effects of retinal/non-retinal, amber/blue light, and similar/dissimilar odors were also confirmed in GraphPad Prism using Welch’s t-test (not assuming equal SD) and by a non-parametric Mann-Whitney test.

## Acknowledgements

We thank B. Gerber for odor containers, A. Cardona for access and training on CATMAID, T. Awasaki, S. Certel, K. Ito, M Landgraf, T. Lee, C.H.-Lee, B Pfeiffer, M. Ramaswami, K. Scott, J Truman, and the Bloomington Drosophila Stock Center (BDSC) for numerous fly stocks, the Cambridge Genetics Department Fly Facility for stocks and microinjections, and the Developmental Studies Hybridoma Bank (DSHB), University of Iowa, for antibodies. We thank M Morgan for help in building optogenetic illumination apparatus.

JYHW was supported by a Medical Research Council studentship and MM was supported by a UK Genetics Society “Genes and Development” summer scholarship and an award from the Bedford Fund, King’s College Cambridge. TB was funded by a BBSRC Doctoral Training Program summer placement grant. This work was supported by BBSRC Grants BB/I022651/1 and BB/N007948/1 and an Isaac Newton Trust award to LMM-N and CJO’K.

## Author contributions

Experimental design: LMMN, CJO’K

Performed experiments: JYHW, BAW, TB, MM, ADM, LMMN, SWZ

Data analysis: LMMN, BAW, JYHW, CJO’K, MM, ADM,

Paper drafting and writing: LMMN

