## Supplemental Figures, Methods, Table for "Octopaminergic neurons have multiple targets in *Drosophila* larval mushroom body calyx and regulate behavioral odor discrimination"

### Supplementary Figure Legends

**Supplementary Fig. 1. *LexA* or *GAL-4* GRASP controls.** To verify that only puncta of reconstituted GFP were detected in GRASP experiments, a line carrying GRASP constructs *UAS-CD4::spGFP1-10* and *LexAop-CD4::spGFP11*, as used in Fig. 3, was crossed to single *LexA* or *GAL4* insertions as shown, and GRASP signals were detected in the larval progeny. Panels show sections of calyces of $3^{rd}$ instar larvae, labeled with monoclonal rat anti-GFP, anti-DLG, and anti-octopamine. **A.** Top row: *Tdc2-LexA(II)*, bottom row *Mef2-GAL4*. Intensity levels for image processing were adjusted to the same levels as Fig 3C. Note the complete absence of GFP puncta. **B.** Top row is *Tdc2(III)*; bottom row is *NP225-GAL4*. Note the complete absence of GFP puncta for *Tdc2-LexA(III)*; only an occasional smear over glomeruli was observed when maximum intensity was adjusted to the same levels as Fig. 3B (*NP225-GAL4* + *Tdc2-LexA* (III)). GFP signal was completely absent from *NP225-GAL4* GRASP when intensity levels were adjusted to the same values as Fig. 3B. Scale bar is 10 μm.

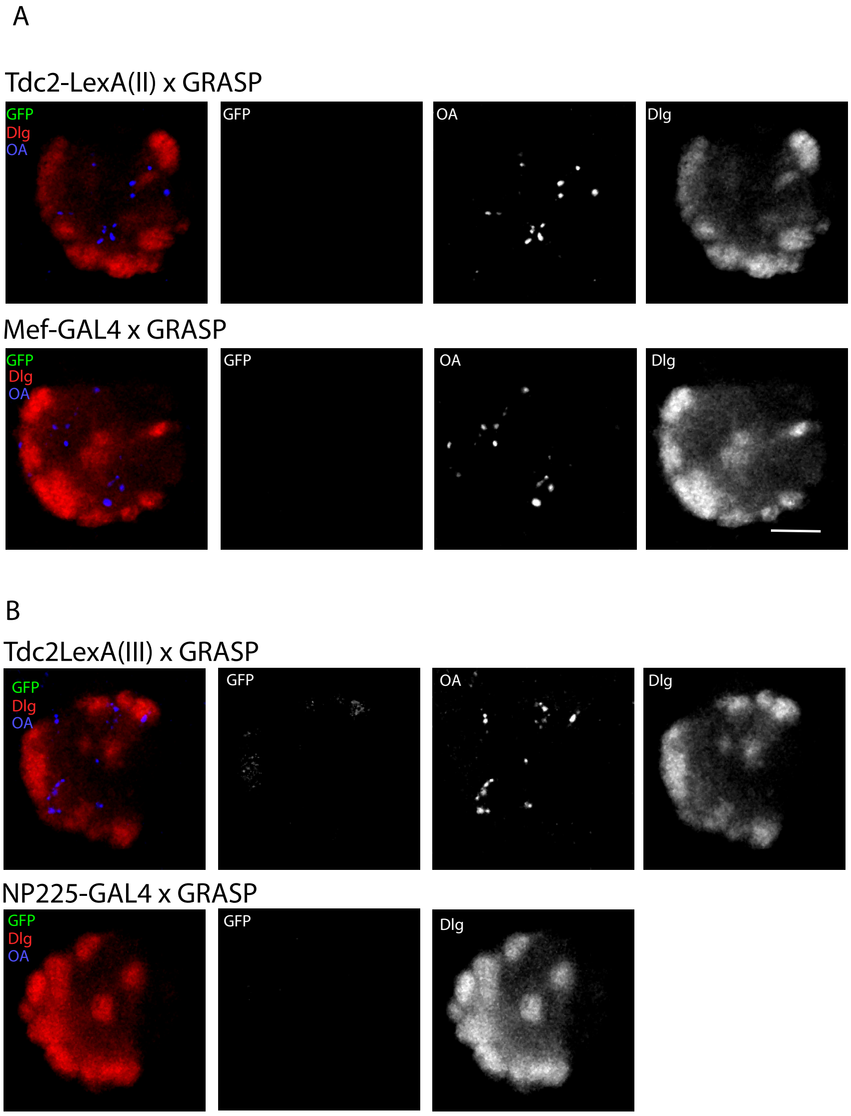

**Supplementary Fig. 2. PCR verification of *MI12417(Oamb)* insertion in the *Oamb* gene. (A)** Primers designed against 5' (*MI12417-5F/MiMIC-5R*) and 3' (*MiMIC-3F/MI12417-3R*) *MiMIC* insertion flanking ends were used to validate the *MI12417* insertion in the *Oamb* gene. **(B)** PCR results for *MI12417* MiMIC insertion. PCR products were detected for *MI12417* 5' and 3' flanking ends using *MI12417* DNA template, but not for the negative control template (denoted as +). A PCR product was detected using primers against the *Oamb* genomic flanking sequences (*MI12417-5F/MI12417-3R)* for the negative control lacking *MI12417* insertion, as well as for *MI12417* due to heterozygosity of the insertion. Abbreviations: MiL/MiR, *MiMIC* insertion ends Left/Right. SA, Splice Acceptor Site.

**A**

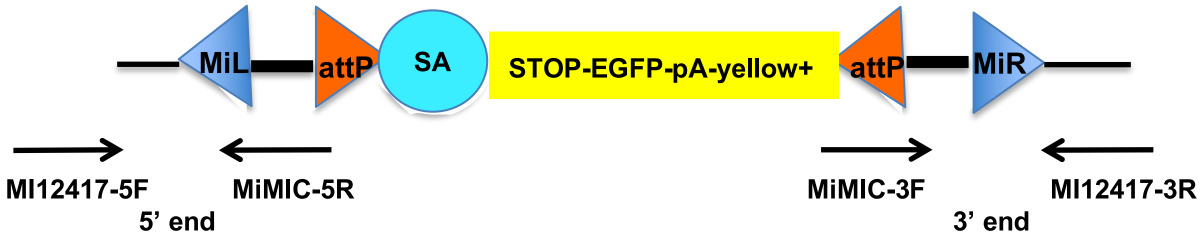


**B**

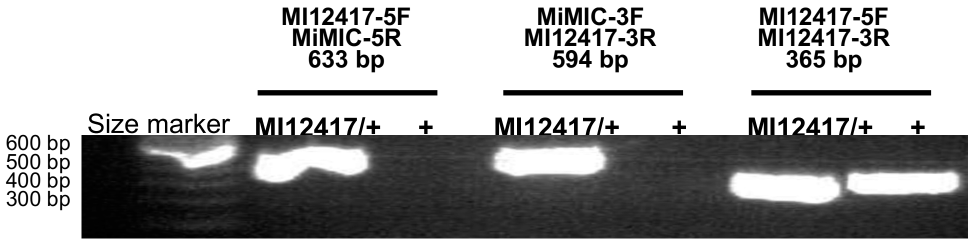

**Supplementary Fig. 3. Alignment of *MI12417* 5' flanking PCR products to MiMIC and Oamb genomic sequences. (A)** Sequenced 5' *MI12417* PCR product. Alignment to MiMIC sequences is indicated in yellow, alignment to *Oamb* in grey. Overlapping alignment at the MiMIC insertion site (TA) indicated in red. **(B-C)** Alignment of *MI12417* 5' PCR products to MiMIC **(B)** and *Drosophila melanogaster* **(C)** sequences using Nucleotide BLAST (Altschul et al., 1990).

### A. Sequence of *MI12417* 5' flanking end

```
NTCTTTATTCGCATAATATAAATCACATGGCTCAAAGAAGAAGTCACGCATCATAATTAG
CATTATATTCCAAGTTGGGCAAAAACAAGAAATCGATAAACGCTGAGGAAGCACATCAAA
CAAATCGGCTTAGATAAAATTGAGGTTGGGATTAAATAGGAGCGGGATTCCAGCCAGAAA
ATGGCAGACATGAAAGCGAGCCATCGGCTAAAACGAAATAAAATATACGAGCCCCAACCA
CTATTAATTCGAACAGCATGTTTTTTTGCAGTGCGCAATGTTTAACACACTATATTATCA
ATACTACTAAAGATAACACATACCAATGCATTTCGTCTCAAAGAGAATTTTATTCTCTTC
ACGACGAAAAAAAAAGTTTTGCTCTATTTCCAACAACAACAAAAATATGAGTAATTTATT
CAAACGGTTTGCTTAAGAGATAAGAAAAAAGTGACCACTATTAATTCGAACGCGGCGTAA
GCTACTAAATCTCTCANGAA
```

### B. *MI12417* 5' flanking end alignment with *Oamb*

```
>AE014297.3 Drosophila melanogaster chromosome 3R      Length=32079331
Features in this part of subject sequence:
   octopamine receptor in mushroom bodies, isoform D
   octopamine receptor in mushroom bodies, isoform G

Strand=Plus/Minus
Query  71        CAAGTTGGGCAAAAACAAGAAATCGATAAACGCTGAGGAAGCACATCAAACAAATCGGCT  130
                 ||||||||||||||||||||||||||||||||||||||||||||||||||||||||||||
Sbjct  20697214  CAAGTTGGGCAAAAACAAGAAATCGATAAACGCTGAGGAAGCACATCAAACAAATCGGCT  20697155

Query  131       TAGATAAAATTGAGGTTGGGATTAAATAGGAGCGGGATTCCAGCCAGAAAATGGCAGACA  190
                 ||||||||||||||||||||||||||||||||||||||||||||||||||||||||||||
Sbjct  20697154  TAGATAAAATTGAGGTTGGGATTAAATAGGAGCGGGATTCCAGCCAGAAAATGGCAGACA  20697095

Query  191       TGAAAGCGAGCCATCGGCTAAAACGAAATAAAATATA  227
                 |||||||||||||||||||||||||||||||||||||
Sbjct  20697094  TGAAAGCGAGCCATCGGCTAAAACGAAATAAAATATA  20697058
```

### C. *MI12417* 5' flanking end alignment with MiMIC sequences

```
>GU370067.1 Synthetic construct MIMIC transposable element, complete sequence   Length=7267

Strand=Plus/Plus
Query  226  TACGAGCCCCAACCACTATTAATTCGAACAGCATGtttttttGCAGTGCGCAATGTTTAA  285
            ||||||||||||||||||||||||||||||||||||||||||||||||||||||||||||
Sbjct  102  TACGAGCCCCAACCACTATTAATTCGAACAGCATGTTTTTTTGCAGTGCGCAATGTTTAA  161

Query  286  CACACTATATTATCAATACTACTAAAGATAACACATACCAATGCATTTCGTCTCAAAGAG  345
            ||||||||||||||||||||||||||||||||||||||||||||||||||||||||||||
Sbjct  162  CACACTATATTATCAATACTACTAAAGATAACACATACCAATGCATTTCGTCTCAAAGAG  221

Query  346  AATTTTATTCTCTTCACGACGaaaaaaaaaGTTTTGCTCTATTTCCAACAACAACAAAAA  405
            ||||||||||||||||||||||||||||||||||||||||||||||||||||||||||||
Sbjct  222  AATTTTATTCTCTTCACGACGAAAAAAAAAGTTTTGCTCTATTTCCAACAACAACAAAAA  281

Query  406  TATGAGTAATTTATTCAAACGGTTTGCTTAAGAGATAAGAAAAAAGTGACCACTATTAAT  465
            ||||||||||||||||||||||||||||||||||||||||||||||||||||||||||||
Sbjct  282  TATGAGTAATTTATTCAAACGGTTTGCTTAAGAGATAAGAAAAAAGTGACCACTATTAAT  341
```

**Supplementary Fig. 4. Alignment of 3' flanking PCR products from *MI12417* to MiMIC and Oamb genomic sequences (A)** Sequenced 3' *MI12417* PCR product. Alignment to MiMIC sequences is indicated in yellow, alignment to *Oamb* sequences in grey. Overlapping alignment at the MiMIC insertion site (TA) indicated in red. **(B-C)** Alignment of *MI12417* 3' PCR products to MiMIC **(B)** and *Drosophila melanogaster* **(C)** sequences using Nucleotide BLAST (Altschul et al., 1990).

### A. Sequence of *MI12417* 3' flanking end

```
GCGGGAGTCGCGACTACGCCCCCACTGAGAGACTCAAAGGTTTACCCCAGTTGGGGCACT
ACTCCCGAAAACCGCTTCTGACCTGGGCCGCGGGGGAAATTAATTAAATTATTGTTTTAA
GTATGATAGTAAATCACATTACGCCGCGTTCGAATTAATAGTGGTCACTTTTTTCTTATC
TCTTAAGCAAACCGTTTGAATAAATTACTCATATTTTTGTTGTTGTTGGAAATAGAGCAA
AACTTTTTTTTTCGTCGTGAAGAGAATAAAATTCTCTTTGAGACGAAATGCATTGGTATG
TGTTATCTTTAGTAGTATTGATAATATAGTGTGTTAAACATTGCGCACTGCAAAAAAAAC
ATGCTGTTCGAATTAATAGTGGTTGGGGCTCGTAATATGTCTTCCCTGTTAGCATGTTCT
GTTTGCAATTTTCTATTTTCTTAGGTTTTTGTCGTTTCAGGCCTCACTGGTCCCCAAAGA
CTCTGTGGGGCCGGATAATCGGCTTTGTTCTGACAGCCGTTTTTTTGCTGGGCTGAATGT
TTAACACACTGGACCATCAGTTTGACTCAGGACTCACGGACCAGTGGCTCCTCCTCTCAA
AGAGAGTTTTATTCTCCTCGCCACAGCAAGGAAGGCCCTGCACGATGTCGAACAGGACCC
CGTGTGGCTCATCTCTTGCTTCTGTGGGAGGCAAGTCTAACCCAGTGTGACCTCCATGAA
GTCGAGAACAAGTAACTCAATCTCCCATCCCACCCTTATGCTGCGCCAGACCCGTGAGGA
GCCCACCTCCGGTGGACAC
```

### B. *MI12417* 3' flanking end alignment with MiMIC sequences

```
>AE014297.3 Drosophila melanogaster chromosome 3R      Length=32079331
Features in this part of subject sequence:
   octopamine receptor in mushroom bodies, isoform D
   octopamine receptor in mushroom bodies, isoform G

>GU370067.1 Synthetic construct MIMIC transposable element, complete sequence   Length=7267
Strand=Plus/Plus
Query  252   tCGTCGTGAAGAGAATAAAATTCTCTTTGAGACGAAATGCATTGGTATGTGTTATCTTTA  311
             ||||||||||||||||||||||||||||||||||||||||||||||||||||||||||||
Sbjct  7062  TCGTCGTGAAGAGAATAAAATTCTCTTTGAGACGAAATGCATTGGTATGTGTTATCTTTA  7121

Query  312   GTAGTATTGATAATATAGTGTGTTAAACATTGCGCACTGCaaaaaaaaCATGCTGTTCGA  371
             ||||||||||||||||||||||||||||||||||||||||||||||||||||||||||||
Sbjct  7122  GTAGTATTGATAATATAGTGTGTTAAACATTGCGCACTGCAAAAAAAACATGCTGTTCGA  7181

Query  372   ATTAATAGTGGTTGGGGCTCGTA  394
             |||||||||||||||||||||||
Sbjct  7182  ATTAATAGTGGTTGGGGCTCGTA  7204
```

### C. *MI12417* 3' flanking end alignment with *Oamb* sequences

```
>AE014297.3 Drosophila melanogaster chromosome 3R     Length=32079331
Features in this part of subject sequence:
   octopamine receptor in mushroom bodies, isoform D
   octopamine receptor in mushroom bodies, isoform G

Strand=Plus/Minus
Query  393       TAATATGTCTTCCCTGTTAGCATGTTCTGTTTGCAATTTTCTATTTTCTTAGGTTTTTGT  452
                 ||||||||||||||||||||||||||||||||||||||||||||||||||||||||||||
Sbjct  20697059  TAATATGTCTTCCCTGTTAGCATGTTCTGTTTGCAATTTTCTATTTTCTTAGGTTTTTGT  20697000

Query  453       CGTTTCAGGCCTCACTGGTCCCCAAAGACTCTGTGGGGCCGGATAATC  500
                 ||||||||||||||||||||||||||||| |||||| | || ||||||
Sbjct  20696999  CGTTTCAGGCCTCACTGGTCCCCAAAGACGCTGTGGTGGCG-ATAATC  20696953
```

**Supplementary Fig. 5. *MI12417* is inserted in coding intron 3 of *Oamb*. (A)** Extract of TBLASTN query of Oamb-B against the *Drosophila* genome assembly, showing an intron at coordinates 20693849–20698333, inserted at amino-acid residue 338 of Oamb-B. **(B)**. Extract of BLASTN search of *Oamb-B* RNA, showing a splice site between nucleotides 2007 and 2008. **(C)** Alignment of Oamb-B protein and Oamb-B RNA sequences around the third intron, highlighting residue G338 and the AGGG splice site in yellow. **(D)** Map of *MI12417* insertion (3R:20,697,059) relative to *Oamb* gene and transcripts (Adapted from GBrowse, www.flybase.org).

**A**

```
Oamb-B:       284 PWKCELTNDRGYVLYSALGSFYIPMFVMLFFYWRIYRAAVRTTRAINQGFKTTKG 338
                  PWKCELTNDRGYVLYSALGSFYIPMFVMLFFYWRIYRAAVRTTRAINQGFKTTKG
Genome: 20698497 PWKCELTNDRGYVLYSALGSFYIPMFVMLFFYWRIYRAAVRTTRAINQGFKTTKG 20698333

Oamb-B:       338 GSPRESGNNRVDESQLILRIHRGRPCSTPQRTPLSVHSMSSTLSVNSNGGGGGAVASGLG 397
                  GSPRESGNNRVDESQLILRIHRGRPCSTPQRTPLSVHSMSSTLSVNSNGGGGGAVASGLG
Genome: 20693849 GSPRESGNNRVDESQLILRIHRGRPCSTPQRTPLSVHSMSSTLSVNSNGGGGGAVASGLG 20693670
```

**B**

```
Oamb-B:      1964 TGAGAACGACGAGAGCCATCAACCAGGGCTTCAAGACCACCAAG 2007
                  ||||||||||||||||||||||||||||||||||||||||||||
Genome: 20698379 TGAGAACGACGAGAGCCATCAACCAGGGCTTCAAGACCACCAAG 20698332

Oamb-B:      2008 GGCAGTCCCCGCGAGTCGGGCAACAATCGAGTGGACGAGTCCCAGCTCATATTGCGCATT 2067
                  ||||||||||||||||||||||||||||||||||||||||||||||||||||||||||||
Genome: 20693849 GGCAGTCCCCGCGAGTCGGGCAACAATCGAGTGGACGAGTCCCAGCTCATATTGCGCATT 20693790
```

**C**

```
                              Coding intron 3; 3R:20,697,059
                              Exon 4>3R:20,693,848-20,692,947
    ATCAACCAGGGCTTCAAGACCACCAAGGGCAGTCCCCGCGAGTCGGGCAACAATCGAGTG
329 -I--N--Q--G--F--K--T--T--K--G--S--P--R--E--S--G--N--N--R--V-
```

**D**

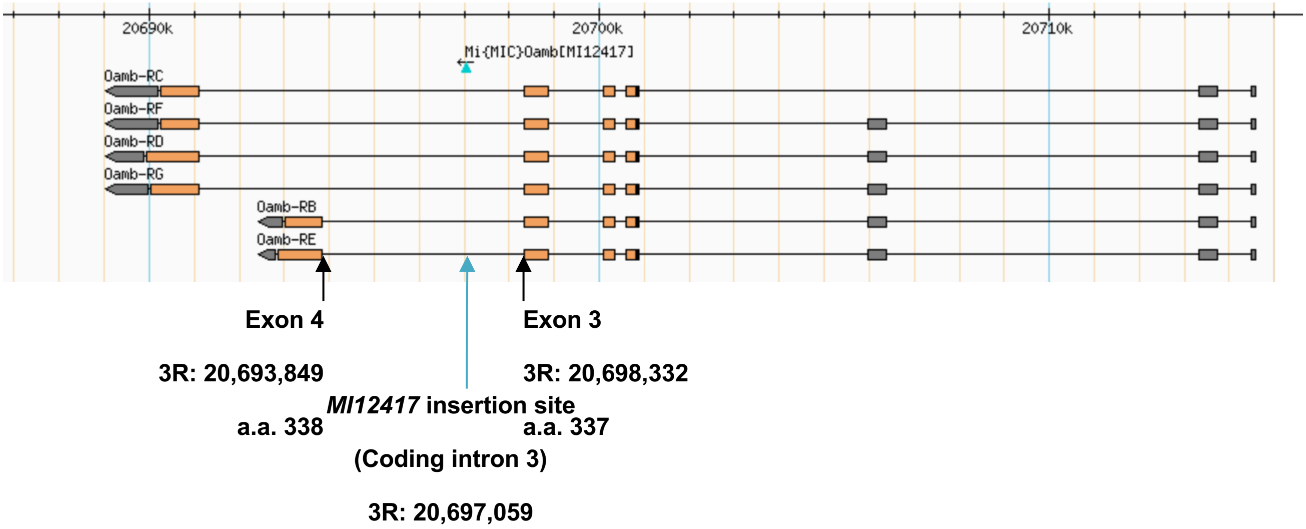

**Supplemmentary Fig. 6.** TMHMM predictions of transmembrane domains for Oamb-PB and PC isotypes. The third intracellular loop, where residue G338 is found, is highlighted in yellow.

#### A. Oamb-PB TMHMM Predictions

| | | | | |
|---|---|---|---|---|
| WEBSEQUENCE | TMHMM2.0 | outside | 1 | 21 |
| WEBSEQUENCE | TMHMM2.0 | TMhelix | 22 | 44 |
| WEBSEQUENCE | TMHMM2.0 | inside | 45 | 56 |
| WEBSEQUENCE | TMHMM2.0 | TMhelix | 57 | 79 |
| WEBSEQUENCE | TMHMM2.0 | outside | 80 | 93 |
| WEBSEQUENCE | TMHMM2.0 | TMhelix | 94 | 116 |
| WEBSEQUENCE | TMHMM2.0 | inside | 117 | 136 |
| WEBSEQUENCE | TMHMM2.0 | TMhelix | 137 | 159 |
| WEBSEQUENCE | TMHMM2.0 | outside | 160 | 293 |
| WEBSEQUENCE | TMHMM2.0 | TMhelix | 294 | 316 |
| WEBSEQUENCE | TMHMM2.0 | inside | 317 | 523 |
| WEBSEQUENCE | TMHMM2.0 | TMhelix | 524 | 546 |
| WEBSEQUENCE | TMHMM2.0 | outside | 547 | 555 |
| WEBSEQUENCE | TMHMM2.0 | TMhelix | 556 | 578 |
| WEBSEQUENCE | TMHMM2.0 | inside | 579 | 637 |

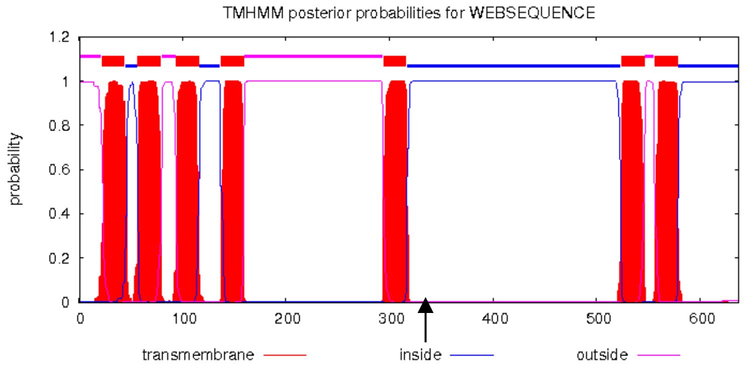


#### B. Oamb-PC TMHMM Predictions

| | | | | |
|---|---|---|---|---|
| WEBSEQUENCE | TMHMM2.0 | outside | 1 | 21 |
| WEBSEQUENCE | TMHMM2.0 | TMhelix | 22 | 44 |
| WEBSEQUENCE | TMHMM2.0 | inside | 45 | 56 |
| WEBSEQUENCE | TMHMM2.0 | TMhelix | 57 | 79 |
| WEBSEQUENCE | TMHMM2.0 | outside | 80 | 93 |
| WEBSEQUENCE | TMHMM2.0 | TMhelix | 94 | 116 |
| WEBSEQUENCE | TMHMM2.0 | inside | 117 | 136 |
| WEBSEQUENCE | TMHMM2.0 | TMhelix | 137 | 159 |
| WEBSEQUENCE | TMHMM2.0 | outside | 160 | 293 |
| WEBSEQUENCE | TMHMM2.0 | TMhelix | 294 | 316 |
| WEBSEQUENCE | TMHMM2.0 | inside | 317 | 520 |
| WEBSEQUENCE | TMHMM2.0 | TMhelix | 521 | 543 |
| WEBSEQUENCE | TMHMM2.0 | outside | 544 | 552 |
| WEBSEQUENCE | TMHMM2.0 | TMhelix | 553 | 575 |
| WEBSEQUENCE | TMHMM2.0 | inside | 576 | 645 |

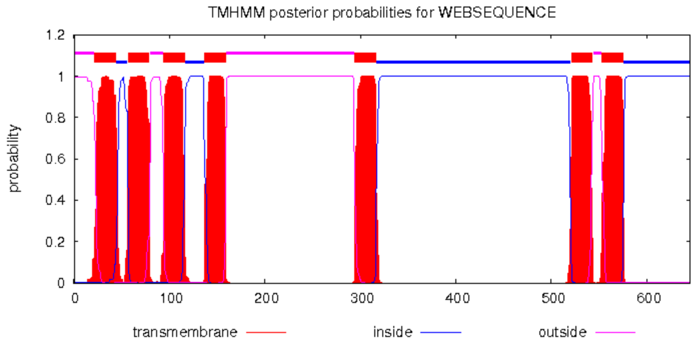

**Supplementary Fig. 7. *MI12417* insertion location relative to *Oamb* cDNA and protein sequences.** cDNA and protein sequences were taken from the ENSEMBL database (Aken et al., 2016; http://www.ensembl.org) for representative *Oamb* transcript protein isotype B. The beginning and range of each exon, obtained from the ENSEMBL database, is annotated in yellow. Transmembrane domains are indicated in green. MiMIC insertion location site, listed on the Gene Disruption Project database, is annotated in light blue. Abbreviations: TM, transmembrane domain; a.a., amino acid

```
3R:20,700,845
Exon 1>3R:20,700,845-20,700,594
    1  ATGAATGAAACAGAGTGCGAGGAT
    1  -M--N--E--T--E--C--E--D-
                                                   TM I: a.a.22-44
   25 CTCATCAAATCTGTGAAATGGACGGAACCAGCCAATCTGATCTCCCTGGCCGTACTCGAG
    9 -L--I--K--S--V--K--W--T--E--P--A--N--L--I--S--L--A--V--L--E-

   85 TTCATCAACGTTCTGGTCATCGGTGGCAACTGCCTCGTGATTGCCGCCGTCTTCTGTTCG
   29 -F--I--N--V--L--V--I--G--G--N--C--L--V--I--A--A--V--F--C--S-

                               TM II: a.a.57-79
  145 AATAAGTTGAGGAGTGTGACGAACTTCTTTATTGTCAACCTAGCTGTGGCCGATCTTCTG
   49 -N--K--L--R--S--V--T--N--F--F--I--V--N--L--A--V--A--D--L--L-

                                                         Exon 2>3R:20,700,331
  205 GTGGGTTTGGCCGTCCTACCCTTCTCAGCCACCTGGGAAGTCTTCAAGGTTTGGATATTC
   69 -V--G--L--A--V--L--P--F--S--A--T--W--E--V--F--K--V--W--I--F-

     -20,700,095      TM III: a.a.94-116
  265 GGCGATCTCTGGTGCCGCATTTGGCTGGCTGTCGATGTCTGGATGTGCACGGCATCGATC
   89 -G--D--L--W--C--R--I--W--L--A--V--D--V--W--M--C--T--A--S--I-

  325 CTGAATCTGTGTGCCATATCACTGGACCGCTATGTGGCGGTCACACGACCCGTCACCTAC
  109 -L--N--L--C--A--I--S--L--D--R--Y--V--A--V--T--R--P--V--T--Y-

                               TM IV: a.a.137-159
  385 CCAAGCATAATGTCCACGAAGAAGGCCAAGTCCTTAATCGCCGGCATTTGGGTACTCTCA
  129 -P--S--I--M--S--T--K--K--A--K--S--L--I--A--G--I--W--V--L--S-

                                                      Exon 3>3R:20,698,857-
  445 TTTTTATTTGCTTTCCGCCGCTAGTCGGCTGGAAGGATCAAAAGGCGGTTATACAGCCG
  149 -F--F--I--C--F--P--P--L--V--G--W--K--D--Q--K--A--V--I--Q--P-

      20,698,335
  505 ACCTATCCAAAGGGAAACCATACGCTTTACTACACCACCACGATGTCAAGCTCGGAGGAT
  169 -T--Y--P--K--G--N--H--T--L--Y--Y--T--T--T--M--S--S--S--E--D-

  565 GGTCAACTAGGGTTAGATAGCATTAAGGACCAGGGCGAGGCATCCTTGCCTCCATCCCCG
  189 -G--Q--L--G--L--D--S--I--K--D--Q--G--E--A--S--L--P--P--S--P-

  625 CCCCATATCGGCAACGGCAACGCCTACAATCCCTACGATCCCGGTTTCGCACCCATCGAT
  209 -P--H--I--G--N--G--N--A--Y--N--P--Y--D--P--G--F--A--P--I--D-

  685 GGATCCGCGGAGATTCGGATTGCGGCCATTGACTCGACCAGTACTTCAACAACCGCAACC
  229 -G--S--A--E--I--R--I--A--A--I--D--S--T--S--T--S--T--T--A--T-

  745 ACCACGACGACAGCGTCCAGCTCGAGCACCACGGAAACGGAAATGGACCTCGATCTACTG
  249 -T--T--T--T--A--S--S--S--S--T--T--E--T--E--M--D--L--D--L--L-

  805 AACGCACCGCCGCAGAACAGACCCCAAACAATTTCCGGCAGTTGTCCGTGGAAGTGCGAG
  269 -N--A--P--P--Q--N--R--P--Q--T--I--S--G--S--C--P--W--K--C--E-
```

```
                     TM V: a.a.294-316
 865 CTGACCAACGATCGGGGTTATGTCCTGTACTCCGCCCTGGGCTCATTCTATATACCCATG
 289 -L--T--N--D--R--G--Y--V--L--Y--S--A--L--G--S--F--Y--I--P--M-

 925 TTCGTGATGCTCTTCTTCTACTGGCGCATCTACCGGGCTGCCGTGAGAACGACGAGAGCC
 309 -F--V--M--L--F--F--Y--W--R--I--Y--R--A--A--V--R--T--T--R--A-

                                  MI12417> Coding intron 3; 3R:20,697,059
                                           MI12417> Intracellular: a.a.338
                                  Exon 4>3R:20,693,848-20,692,947
 985 ATCAACCAGGGCTTCAAGACCACCAAGGGCAGTCCCCGCGAGTCGGGCAACAATCGAGTG
 329 -I--N--Q--G--F--K--T--T--K--G--S--P--R--E--S--G--N--N--R--V-

1045 GACGAGTCCCAGCTCATATTGCGCATTCACCGAGGAAGACCTTGCTCCACCCCCCAGCGC
 349 -D--E--S--Q--L--I--L--R--I--H--R--G--R--P--C--S--T--P--Q--R-

1105 ACGCCCCTCTCGGTGCACTCAATGTCCTCGACTCTCAGCGTGAACAGCAACGGGGGCGGG
 369 -T--P--L--S--V--H--S--M--S--S--T--L--S--V--N--S--N--G--G--G-

1165 GGTGGAGCCGTGGCCTCGGGACTGGGTGCCTCCACCGAGGATCACCTTCAGGGAGGCGCC
 389 -G--G--A--V--A--S--G--L--G--A--S--T--E--D--H--L--Q--G--G--A-

1225 CCCAAGCGGGCCACATCGATGCGCGTCTGCCGACAGCGACACGAGAAGGTGGCCATCAAG
 409 -P--K--R--A--T--S--M--R--V--C--R--Q--R--H--E--K--V--A--I--K-

1285 GTGTCCTTTCCCTCCTCCGAGAATGTCCTCGACGCAGGACAGCAGCCACAGGCATCGCCA
 429 -V--S--F--P--S--S--E--N--V--L--D--A--G--Q--Q--P--Q--A--S--P-

1345 CACTATGCGGTAATCAGTAGCGCCAACGGACGTCGTGCCTCCTTTAAGACGAGCCTCTTC
 449 -H--Y--A--V--I--S--S--A--N--G--R--R--A--S--F--K--T--S--L--F-

1405 GACATTGGCGAGACCACCTTTAATTTGGACGCAGCTGCGTCCGGTCCCGGAGACCTAGAG
 469 -D--I--G--E--T--T--F--N--L--D--A--A--A--S--G--P--G--D--L--E-

1465 ACCGGACTCTCGACCACCTCACTGTCGGCCAAGAAGCGGGCAGGCAAGCGCAGCGCCAAG
 489 -T--G--L--S--T--T--S--L--S--A--K--K--R--A--G--K--R--S--A--K-

                                                TM VI: a.a.524-546
1525 TTTCAGGTGAAGCGGTTCCGAATGGAGACCAAGGCAGCCAAGACGCTGGCCATCATTGTG
 509 -F--Q--V--K--R--F--R--M--E--T--K--A--A--K--T--L--A--I--I--V-

1585 GGCGGCTTCATCGTTTGCTGGCTGCCCTTCTTCACGATGTATCTGATCCGGGCCTTCTGC
 529 -G--G--F--I--V--C--W--L--P--F--F--T--M--Y--L--I--R--A--F--C-

                         TM VII: a.a.556-578
1645 GACCACTGCATTCAGCCGACGGTCTTTTCGGTGCTCTTCTGGCTGGGCTACTGCAACTCG
 549 -D--H--C--I--Q--P--T--V--F--S--V--L--F--W--L--G--Y--C--N--S-

1705 GCCATTAATCCGATGATCTATGCGCTCTTCTCGAATGAGTTTCGCATCGCCTTCAAGCGG
 569 -A--I--N--P--M--I--Y--A--L--F--S--N--E--F--R--I--A--F--K--R-

1765 ATAGTGTGCAGATGCGTCTGCACCCGCAGTGGCTTCCGGGCGTCGGAGAATTTCCAGATG
 589 -I--V--C--R--C--V--C--T--R--S--G--F--R--A--S--E--N--F--Q--M-

1825 ATAGCGGCGCGTGCCCTGATGGCACCGGCAACATTCCACAAGACCATATCCGGATGCTCG
 609 -I--A--A--R--A--L--M--A--P--A--T--F--H--K--T--I--S--G--C--S-

1885 GACGACGGCGAGGGCGTGGACTTCAGCTGA
 629 -D--D--G--E--G--V--D--F--S--*-
```

**Supplementary Table S1.**

**Comparison of synapse numbers between sVUM1 neurons and postsynaptic neurons in the calyx in 1st and 3rd instar larvae.**

The number of synapses between sVUM1 neurons and other neurons in the calyx (KCs, APL, both Odd neurons together) for first instar larva were extracted from the online publicly available resource, Virtual Fly Brain, https://catmaid.virtualflybrain.org/L1 Larval CNS (L1EM) (Licence CC-BY-SA_4.0). First instar data correspond to synaptic contacts judged by EM criteria, corresponding approximately to the numbers of active zones. These were obtained by using the connectivity tool to display all downstream neurons of OAN-a1/ sVUMmd1 or OAN-a2/ sVUMmx1, and exported as a csv file. Numbers indicate the total number of synapses per neuron type with both sVUM1s, in both brain hemispheres, for APL and Odds, extracted from the left and right annotations in the CATMAID database. For KCs, the numbers should be doubled as shown for consistency, since the numbers were manually calculated from the csv file by identifying the numbers of synapses corresponding to single KCs, and these were annotated only on one side of the brain, in the CATMAID database. Calyx sVUM1-PN synapses are not included, since these could not be distinguished in these lists from sVUM1-PN synapses in the AL. A breakdown of synapse numbers for individual neurons in first instar larvae is in Supplementary File 1.

Third instar data were taken from OA-positive GRASP puncta as described in the Results text of this work, and refer to one brain hemisphere only. For the purpose of comparisons, numbers must be doubled as indicated to represent values for both brain halves together. Examples are shown in Fig. 3. Notice almost double numbers of synapses at 3rd instar larval stage for KCs and Odd, while for APL, numbers at first instar are 11.6% of that at 3rd instar.

| | KCs | Odd neurons | APL |
|---|---|---|---|
| **1st instar**<br>KCs: one calyx only<br>Odds and APL: left and right calyx together) | 2 x 49 (n=1) | 67 (n=1) | 13 (n=1) |
| **3rd instar**<br>(single calyx; double this number to estimate left and right calyx together) | 2 x 96 ± 34 (n=3) | 2 x 63 ± 3 (n=3) | 2 x 56 ± 7 (n=4) |

**Supplementary Methods**

Odor balancing. For learning experiments, we needed to determine concentrations of ethyl acetate (EA) or pentyl acetate (PA) that gave a preference index (Pref-I, defined in the main paper) close to zero in naive larvae, so that deviations from this score could be used as a measure of learning after conditioning with an appetitive stimulus.

We carried out initial balancing experiments using Canton-S (CS) to determine the range of dilutions of ethyl acetate (EA) or (PA) that were approximately equally attractive to larvae.

| | **Pref-I at 2 min** | **Pref-I at 5 min** |
|---|---|---|
| EA 1:2000 vs PA 1:200 | -0.14 ± 0.06 (n=16) | -0.41 ± 0.07 (n=16). |
| EA 1:2000 vs PA 1:500 | -0.10 ± 0.04 (n=19) | -0.31 ± 0.05 (n=19). |
| EA 1:2000 vs PA 1:1000 | -0.03 ± 0.07 (n=8) | -0.30 ± 0.09 (n=8). |

Odor conditioning. We carried out fructose conditioning on CS larvae using EA at 1:2000 and PA at 1:500 dilutions, using white light and standard food. We recorded the Performance Index (PI) values for EA conditioning (EA+), and for PA conditioning (PA+) as defined in the main text, and used these to calculate Performance Index as a measure of learning. Counting of larvae was done at 2 min and 5 min in each test.

| | **PI (EA+)** | **PI (PA+)** | **PI (Learning)** |
|---|---|---|---|
| 2 min | 0.27 ± 0.04 (n=18) | -0.27 ± 0.05 (n=18) | 0.27 ± 0.04 (n=18) |
| 5 min | 0.28 ± 0.06 (n=18) | -0.52 ± 0.03 (n=18) | 0.40 ± 0.04 (n=18). |

We next used these concentrations to measure Pref-I values of larvae of the genotype used in Fig. 9 (n=8)

| **Light** | **Odors** | **Naive Pref-I** |
|---|---|---|
| Blue | EA (1:2000) vs PA (1:500) | 0.03 ±0.06 |
| Blue | EA:PA 1:4 vs EA:PA 4:1, | 0.00 ±0.04 |
| Amber | EA;PA 1:4 vs EA:PA 4:1, | 0.11 ±0.03 |

Given the low naive Pref-I values measures with these odor concentrations, we used these for the learning and discrimination experiments reported in the main paper.
